# Integrated multi-omics reveals coordinated *Staphylococcus aureus* metabolic, iron transport, and stress responses to human serum

**DOI:** 10.1101/2025.08.11.669730

**Authors:** Warasinee Mujchariyakul, Calum J. Walsh, Stefano Giulieri, Cameron Cramond, Kim-Anh LêCao, Timothy P. Stinear, Benjamin P. Howden, Romain Guérillot, Abderrahman Hachani

## Abstract

Bloodstream infections caused by *Staphylococcus aureus* remain a leading cause of mortality worldwide. Our understanding of *S. aureus* survival and persistence in human serum, a cell-free fraction of blood hostile for bacteria, is still limited. Here, we applied multivariate data integration methods and network analysis to a multi-omic dataset generated from five clinically prevalent *S. aureus* genotypes exposed to human serum. We observed and then confirmed using isogenic mutants, the significant roles of *gapdhB*, *sucA*, *sirA*, *sstD*, and *perR* in bacterial survival in serum. These data show that metabolic versatility in carbon source usage, iron transport and resistance to oxidative stress are interlinked and central to *S. aureus* fitness in serum, representing potential *S. aureus* vulnerabilities that could be exploited therapeutically.

**IMPORTANCE:** Bloodstream infections caused by *Staphylococcus aureus* are associated with mortality rates of up to 30%. However, the molecular mechanisms that enable this pathogen to survive in human serum, a nutrient-limited and immunologically hostile environment remain poorly understood. By integrating multi-omic data from five clinically relevant *S. aureus* genotypes and validating key signatures using mutants, we identified conserved genetic determinants critical for bacterial survival in serum. Our findings highlight the interconnected roles of carbohydrate metabolic flexibility, iron acquisition, and oxidative stress resistance in shaping *S. aureus* adaptation to serum. This work advances our understanding of microbial strategies to survive in the bloodstream and demonstrates the potential of multi-omic integration to uncover therapeutic vulnerabilities in bacterial pathogens.

## INTRODUCTION

*Staphylococcus aureus* is a significant opportunistic human pathogen (1). As a commensal, *S. aureus* coexists with other members of the human microbiome in up to 30% of the human population, typically colonizing the skin or nares, with the potential to become a lethal pathogen (2). *S. aureus* bloodstream infections are greater than 10 in 100,000 people per year, with the severity of infections compounded by the widespread emergence of resistance to last-line antibiotics such as vancomycin, and mortality rates reaching 20% (3).

During bloodstream infections, professional phagocytes and non-cellular immune components, including antimicrobial peptides (AMPs), immunoglobulins and the complement system, target invasive pathogens for clearance (4, 5). While serum does not contain immune cells, it represents a potent bactericidal environment for blood-invasive bacterial pathogens (6–8); an environment against which *S. aureus* has evolved multiple defense mechanisms (1, 9, 10). The bacterium can evade humoral immunity by directly interfering with antibody functions to prevent opsonization for phagocytosis (11, 12), or by degrading complement components (13). *S. aureus* can inactivate AMPs through proteolytic degradation (14), modification of its surface charge, or remodeling both the organization of its cell wall peptidoglycan and the abundance of cardiolipin in its plasma membrane. The latter two processes also contribute to *S. aureus* resistance to cell wall-targeting antibiotics (15). The host can also impose nutritional immunity by sequestering essential nutrients, primarily trace metals like iron, zinc, and manganese to restrict bacterial growth. Bacterial pathogens have evolved mechanisms to circumvent nutritional immunity (16). For example, *S. aureus* has multiple trace metal-sequestering strategies that also contribute to its survival in blood (17).

Elucidating the metabolic reprogramming of *S. aureus* in response to serum-derived stressors—such as nutrient limitation and acellular immune factors—can uncover fundamental metabolic pathways that facilitate its adaptation, survival, and persistence in the bloodstream. These insights may inform the development of targeted antimicrobial strategies and rationally designed interventions to mitigate invasive and chronic *S. aureus* infections.

We previously reported a comprehensive multi-omics analysis of four clinically significant sepsis-causing bacterial pathogens (*Escherichia coli*, *Klebsiella pneumoniae, S. aureus*, and *Streptococcus pyogenes*) following their exposure to human serum (18). This work revealed a set of conserved adaptive responses across these phylogenetically diverse species.

Notably, all four pathogens exhibited a coordinated upregulation of lipid metabolic pathways that was associated with extensive remodeling of the cell envelope. This adaptation likely reflects a shared osmoprotective strategy, enhancing bacterial survival in the hostile and nutrient-limited environment of human serum. Building on our multi-omics dataset, this study examines the serum response of *Staphylococcus aureus* across five clinically relevant clonal lineages—BPH2760 (ST1), BPH2819 (ST5), BPH2900 (ST22), BPH2947 (ST239), and BPH2986 (ST8). The aim is to identify a conserved set of molecular features that drives species-specific adaptation to serum during septicemia.

With a focused analysis on *S. aureus*, we refined here our analytical workflow. In our previous study (18), we used a late-integration approach, independently analyzing transcriptomic, proteomic, and metabolomic datasets before mapping significant features to metabolic pathways and Gene Ontology terms. While this revealed broad functional patterns, it lacked the resolution to detect coordinated molecular changes across omics layers or to uncover direct cross-omic relationships. To address these limitations, we adopted the DIABLO framework (Data Integration Analysis for Biomarker discovery using Latent cOmponents) from the mixOmics R package, which applies a sparse, multiblock partial least squares (PLS) method, optimized to integrate all omics datasets within a single multivariate model. By selecting a minimal set of features that co-vary across data types and distinguish between serum- and RPMI-exposed conditions, DIABLO enables the identification of cross-omic interactions, reduces the statistical burden of separate analyses, and preserves the integrated biological context of the *S. aureus* serum response (19, 20). We further interrogated these signatures using differential expression and network-based analyses, and identified interconnected metabolic pathways involved in serum adaptation through over-representation analysis (ORA) and Gene Set Enrichment Analysis (GSEA).

Our integrated omic analysis shows that *S. aureus* maintains optimal fitness in serum by undergoing changes in key carbohydrate metabolic processes, iron acquisition, and resistance to oxidative stress pathways. This combined response likely enables *S. aureus* to withstand host stressors such as nutritional immunity and promote its survival in human serum.

## RESULTS

### Coordinated transcriptomic, proteomic, and metabolomic adaptation of *S. aureus* to serum

We performed multi-omics analysis to find the shared molecular responses to human serum of five *S. aureus* clinical isolates (BPH2760, BPH2819, BPH2900, BPH2947, and BPH2986) (**Figure 1A**). Principal Component Analysis (PCA), a dimensionality reduction method, was applied to each omics layer, revealing clear separation between serum- and RPMI-exposed *S. aureus*. The first principal component (PC1) explained 24%, 22%, 27%, and 29%, respectively, of the total variance in the transcriptomic, proteomic, and metabolomic (GC-MS and LC-MS) datasets, capturing the dominant condition-specific signal (**Figure 1B**). We used DIABLO to integrate the datasets into a unified predictive model of *S. aureus* exposure to human serum. DIABLO is a supervised multiblock method based on sparse Partial Least Squares Discriminant Analysis (sPLS-DA) (19, 20). DIABLO uses the PLS framework to model relationships between molecular features (derived from transcriptomic, proteomic or metabolomic datasets) and phenotypic outcomes. By integrating multiple omics datasets (defined as multiblock) and applying feature selection (sparsity), DIABLO highlights the subset of variables with the greatest discriminatory power – in this case, the staphylococcal molecules most strongly differentiating growth in RPMI versus serum (**Figure 1C-D, Figure S1A-C).** The sample plot showed that the first component captured most of the common serum responses across all five *S. aureus* strains, while the second component reflected strain-specific variation (**Figure S1C**). The final DIABLO model used 60 multi-omic features (see methods) that optimally discriminated serum from RPMI exposure on the first component (**Table S1**, **Figure S1D**).

**Figure 1.**
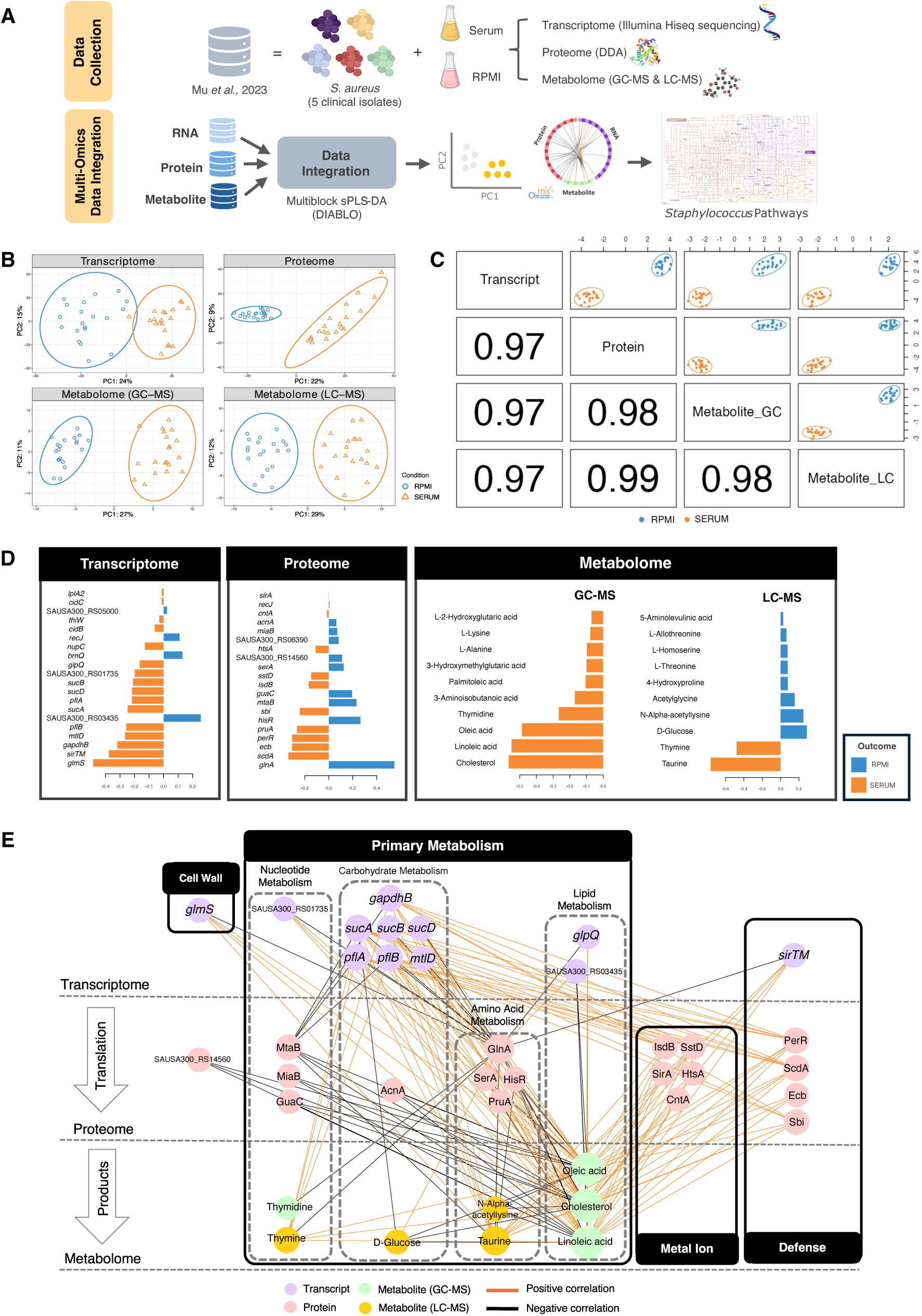
Multi-omics identification of molecular signatures of *S. aureus* adaptation to serum. (A) Schematic workflow of multi-omics analysis to predict *S. aureus* signature molecules in response to human serum; (B) Unsupervised analysis using Principal Component Analysis (PCA). Sample plot from PCA illustrating the consistency of sample clustering across datasets (transcriptomics, proteomics, and metabolomics (GC-MS and LC-MS)), with samples labelled based on media conditions (SERUM and RPMI); (C) Supervised analysis using multiblock sPLS-DA. Diagnostic plot (sample scatter plot), according to media conditions (SERUM and RPMI). The numbers in each block below the diagonal represent correlation coefficients between the first components of each dataset; (D) Loading plot depicting features selected as optimally discriminatory between conditions by PLS-DA from the first component in each dataset; (E) Correlation network visualizing pairwise correlations (> |0.9|) between variables (transcripts, proteins, metabolites). Edge connections represent correlations, with edge colors indicating positive (orange) or negative (black) correlations. Node sizes are proportional to the number of interactions, and node colors represent data types: purple (transcripts), pink (proteins), green (GC-MS metabolites), and yellow (LC-MS metabolites).

A coordinated serum response signature was indicated by highly correlated latent components across the omics layers, including transcriptomic, proteomic and metabolomic. The signatures extracted from the omics layers were highly correlated (Pearson’s r > 0.95) and indicated both class separation and balanced correlation between the datasets (**Figure 1C)**. Loading values from the DIABLO model (**Figure 1D**) show the specific contribution of individual variables, with higher absolute values marking greater discriminative power.

To explore consistent patterns between omics data, a similarity matrix was calculated across all components and selected features (**Table S1**) and visualized as a network (**Figure 1E)** (21). The network revealed interactions across transcriptomic, proteomic and metabolomic layers, with features grouped by function and showing both positive and negative correlations and suggesting a coordinated response to serum. The staphylococcal pathways enriched among the discriminant and correlated features identified by DIABLO included cell wall biosynthesis, carbohydrate metabolism, iron transport, and defense against host immunity (**Figure 1D, 1E and Table 1)**. CDS and their products central to these pathways exhibited strong positive correlations: i) *glmS*, pivotal in the cell wall biosynthesis and lipid metabolism (via linoleic and oleic acids, and cholesterol metabolites); ii) *gapdhB* and *sucA*, involved in carbohydrate metabolism, and *sstD*, *scdA*, and *isdB*, contributing to iron uptake; and iii) *perR* and *sbi*, supporting oxidative stress resistance mechanisms and immune evasion, respectively. In contrast, genes involved in nucleotide metabolism (*mtaB*, *miaB*, *guaC*) were negatively correlated with carbohydrate metabolism genes (*gapdhB* and s*ucA*) and lipids (linoleic and oleic acids).

**Table 1.**
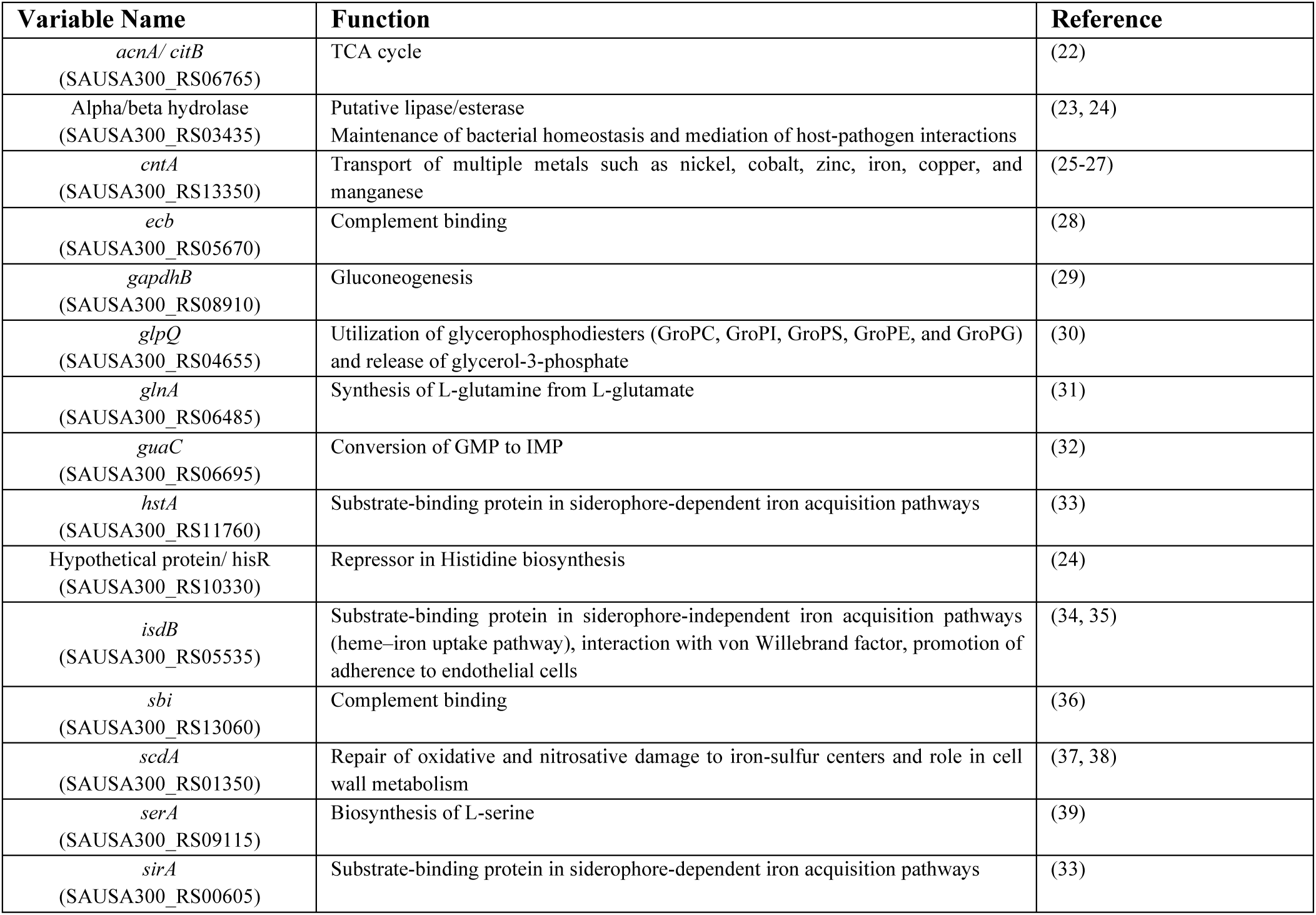
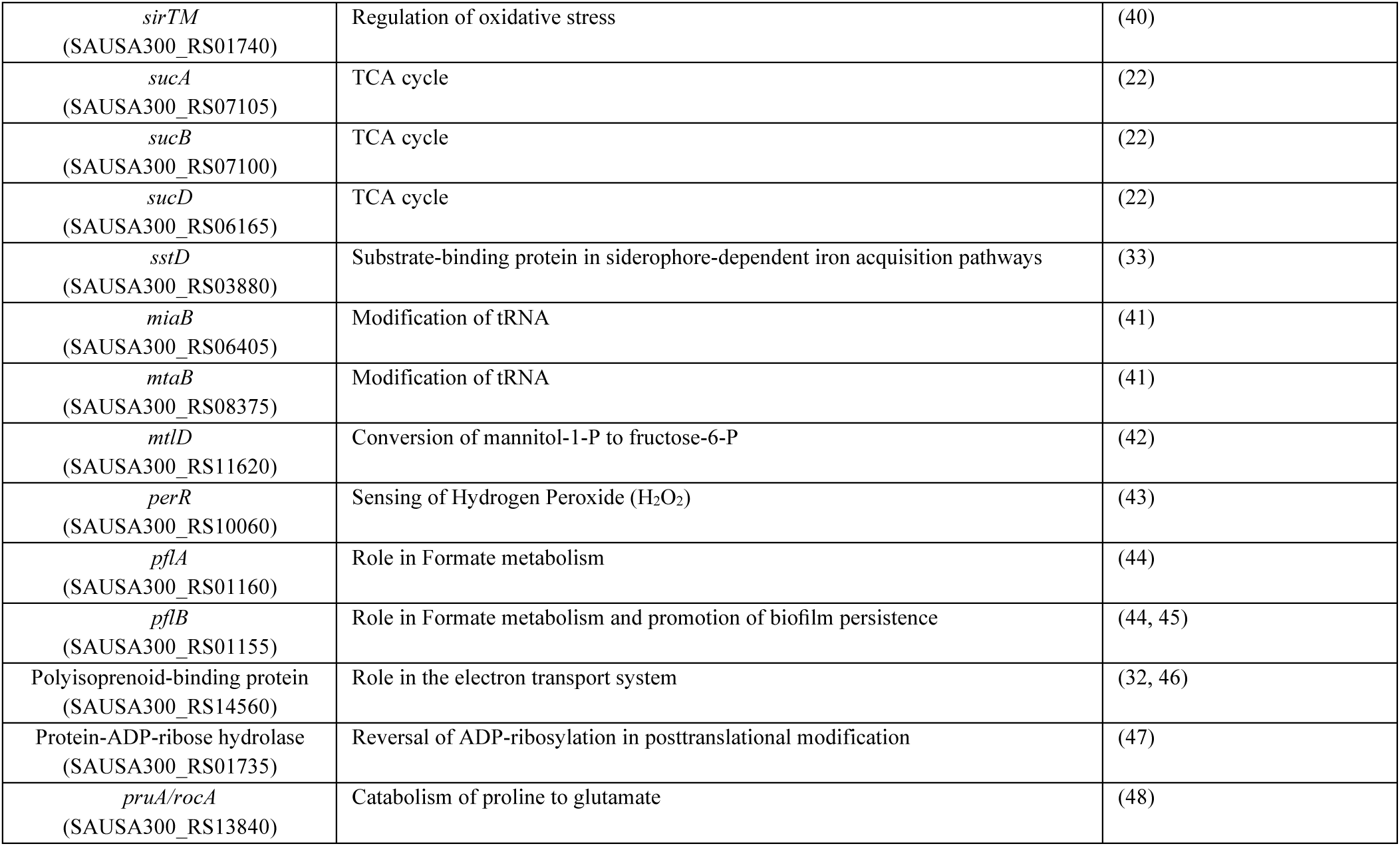
Functions of selected variables shown in Figure 1E.

The observed patterns show that *S. aureus* undergoes substantial metabolic adaptation in response to serum, engaging uptake and utilization of serum-derived amino acids, carbohydrates, fatty acids, nucleotides, and their derivatives. Notably, the analysis revealed a marked increase in the abundance of staphylococcal iron acquisition proteins upon serum exposure. In the host environment, extracellular iron is limited and typically sequestered by high-affinity iron-binding proteins such as transferrin, lactoferrin, and hemoglobin within erythrocytes. The elevated levels of both siderophore-dependent (HtsA, SirA, SstA) and siderophore-independent (IsdB) iron acquisition proteins underscore the critical role of iron sequestration in *S. aureus* survival (**Figure 1D**). The integration of multi-omic data reveals the responses that support metabolic resilience, enabling *S. aureus* to maintain fitness while countering intrinsic serum stressors.

### Network and enrichment analyses reveal *S. aureus* shared and strain-specific responses to serum

Differentially expressed (DE) genes or proteins for each strain were identified and further subjected to an intersection analysis. This analysis showed that at the transcript level, 143 genes were up-regulated, and 65 genes were down-regulated in response to serum. At the proteomic level, there were 21 up-regulated and 17 down-regulated proteins in response to serum (**Figure S2A-B, Table S2**). We performed a Pathway Enrichment Analysis (PEA), including an overrepresentation analysis (ORA) and a gene set enrichment analysis (GSEA), to reduce the complexity of data and find overrepresented pathways elicited upon exposure to serum. ORA is widely used to identify over-represented biological functions in DE gene sets, performing best when handling wide gene expression differences. Genes were filtered using criteria of log2 fold change (log2FC) > 1 and adj-P value < 0.05. To improve the granularity of PEA outputs, a GSEA analysis was also applied to capture pathways where gene expression changes were more subtle but consistently enriched, enabling the detection of significant, but lower-magnitude responses. Using the similarity in gene subsets shared between pathways, we also conducted a network analysis to find interactions among the enriched pathways across the omics layers (**Figure 2A, Figure S3A-J, Table S3**). Consistent with previous DIABLO findings, ORA and GSEA identified four distinct functional modules comprising carbohydrate metabolism, ribosome-associated pathways, iron acquisition, and nucleotide metabolism (**Figure 2B-E**).

**Figure 2.**
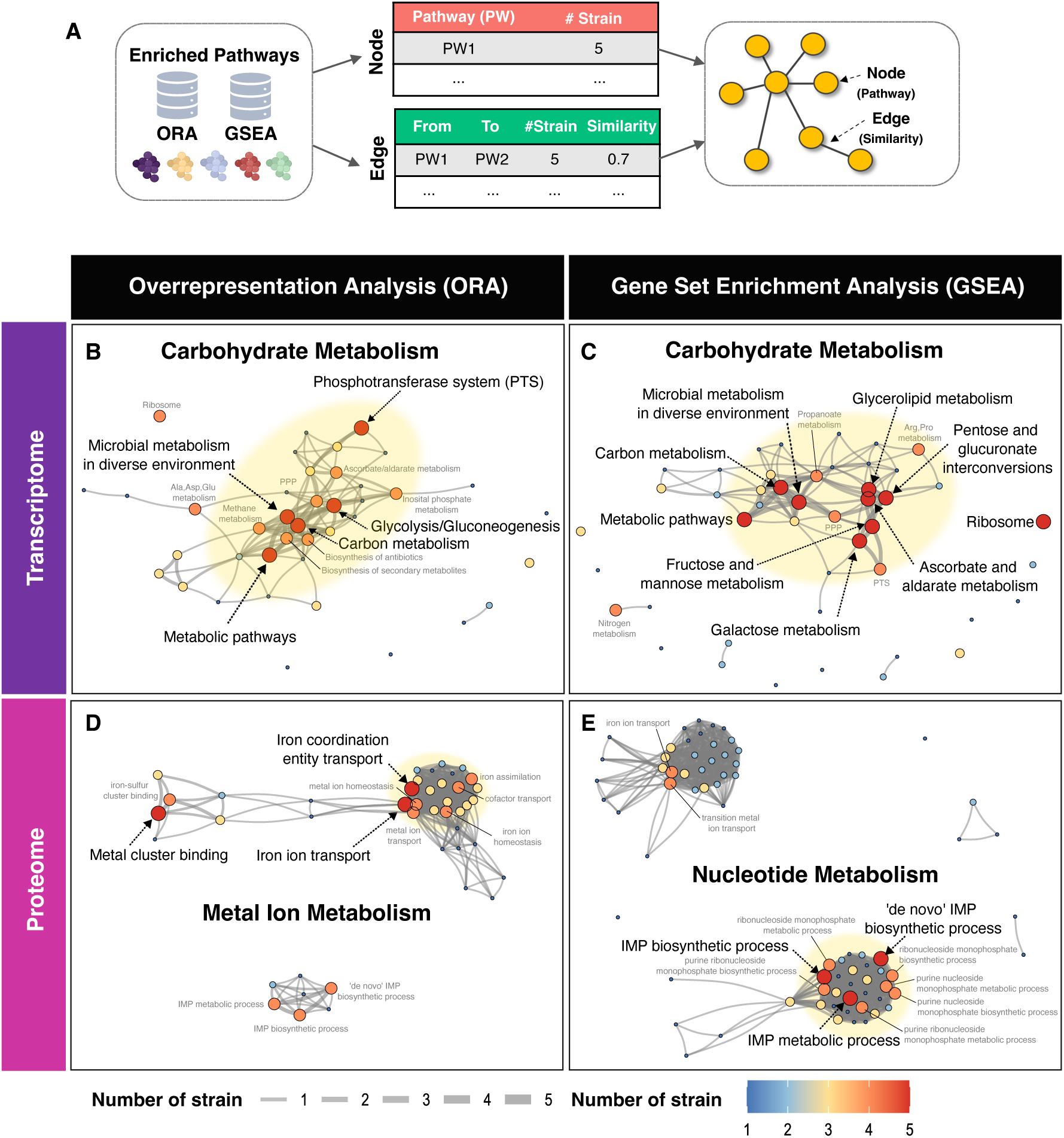
Clustering of transcriptomics and proteomics data into function-related modules. (A) Workflow overview for network analysis of enriched pathways identified from over-representation analysis (ORA) and gene set enrichment analysis (GSEA), with nodes representing pathways and edges representing pairwise similarity (Sim) between enriched terms or pathways (PW), calculated using Jaccard’s similarity coefficient. (B-C) ORA and GSEA of transcriptomic data. (D-E) ORA and GSEA of proteomic data. Dot sizes and colors represent the number of strains sharing the same enriched pathways. Edge widths indicate the number of strains with interactions between pathways, while edge lengths reflect pathway similarity. Yellow shading highlights denote functionally related modules.

The carbohydrate metabolism module was composed of the glycolysis/gluconeogenesis pathway, along with other carbohydrate-processing routes such as phosphotransferase systems (PTS) (**Figure 2B-C**). Upregulated carbohydrate transporter genes, *uhpT*, *mtlA*, *fruA*, *glcC*, *lacE*, SAUSA300_RS01760, and *rbsU,* support the usage of various saccharides, including glucose, fructose, lactose, D-mannitol, and ribose (**Table S2**). Once transported and phosphorylated by PTS, carbohydrates such as glucose and fructose, are processed by glycolysis and converted into ribose-5-phosphate, glyceraldehyde-3-phosphate, and fructose-6-phosphate intermediates. These key metabolites bridge glycolysis to nucleotide and lipid metabolism (49, 50) (further detailed in **Figure 4)**, thereby enhancing the staphylococcal metabolic flexibility in serum. In contrast to other metabolic modules, our network analysis revealed that ribosome-associated pathways were largely isolated, exhibiting no direct connectivity with broader metabolic networks (**Figure 2C, Figure S3A-D**). Transcriptome profiling indicated a suppression of ribosome biogenesis following serum exposure, characterized by the downregulation of genes encoding 30S and 50S ribosomal subunit proteins (**Table S2 and S3)**. Expression of the translational GTPase *typA* (also known as *bipA*), an important regulator of ribosome assembly and translational fidelity under stress conditions (51, 52), was also significantly reduced (**Table S2)**. These findings show that the protein biogenesis is downregulated as part of a stress-adaptive response of *S. aureus* to serum.

Among the proteomic changes induced by serum exposure, iron acquisition and nucleotide metabolism pathways showed the most substantial alterations (**Figure S1C**). Notably, there was a marked increase in the abundance of iron transport proteins, including SirA, FhuC, SstC, SstD, HtsA, IsdB, IsdE, and IsdI. In addition, elevated levels of ScdA and CntA (53) -proteins involved in iron-sulfur cluster binding - alongside the downregulation of *miaB* (54), suggest the activation of iron-sparing strategies by *S. aureus* under serum conditions (**Tables S2** and **S3**). The significant increase in levels of ScdA and CntA, iron cluster binding proteins, concomitant with the downregulation of *miaB*, point to the engagement of staphylococcal iron-sparing strategies during serum exposure (**Table S2** and **S3**). Thus, *S. aureus* coordinates enhanced iron uptake with the regulation of iron incorporation into metalloproteins to efficiently manage iron availability during serum exposure (**Figure 2D**, **Figure S1B** and **S3F**).

Proteins involved in nucleotide metabolism, particularly those associated with the inosine monophosphate (IMP) biosynthetic pathway, were prominently increased across all *S. aureus* isolates (**Figure 2E** and **Figure S3H**). This pathway is central to the synthesis of purine nucleotides such as adenosine monophosphate (AMP) and guanosine monophosphate (GMP), critical molecules for DNA replication, RNA transcription, and cellular proliferation. These shifts underscore the prioritization of iron homeostasis and nucleotide biosynthesis as key adaptive strategies employed by *S. aureus* to support survival and growth in serum.

### Experimental validation of predicted molecular signatures of *S. aureus* responses to human serum

To confirm findings from the multi-omic integration and pathway enrichment analyses, we assessed the fitness of *S. aureus* transposon mutants in genes involved in carbon metabolism, iron acquisition, and oxidative stress resistance, relative to their parental wild-type USA300 JE2 strain, in the presence of serum (**Figures 1E and 3A**). We initially verified the genetic relatedness of the five *S. aureus* bloodstream isolates to the laboratory strain JE2, used for this phenotypic validation. We observed a high conservation of the *S. aureus* core genome (2,222 core genes-representing 71% of JE2 genome- and 903 accessory genes). Strain JE2 was most closely related to the clinical isolate BPH2986, with both belonging to the ST8 USA300 lineage and differing by only 73 single nucleotide polymorphisms (SNPs) (**Figure S4A-D**).

**Figure 3.**
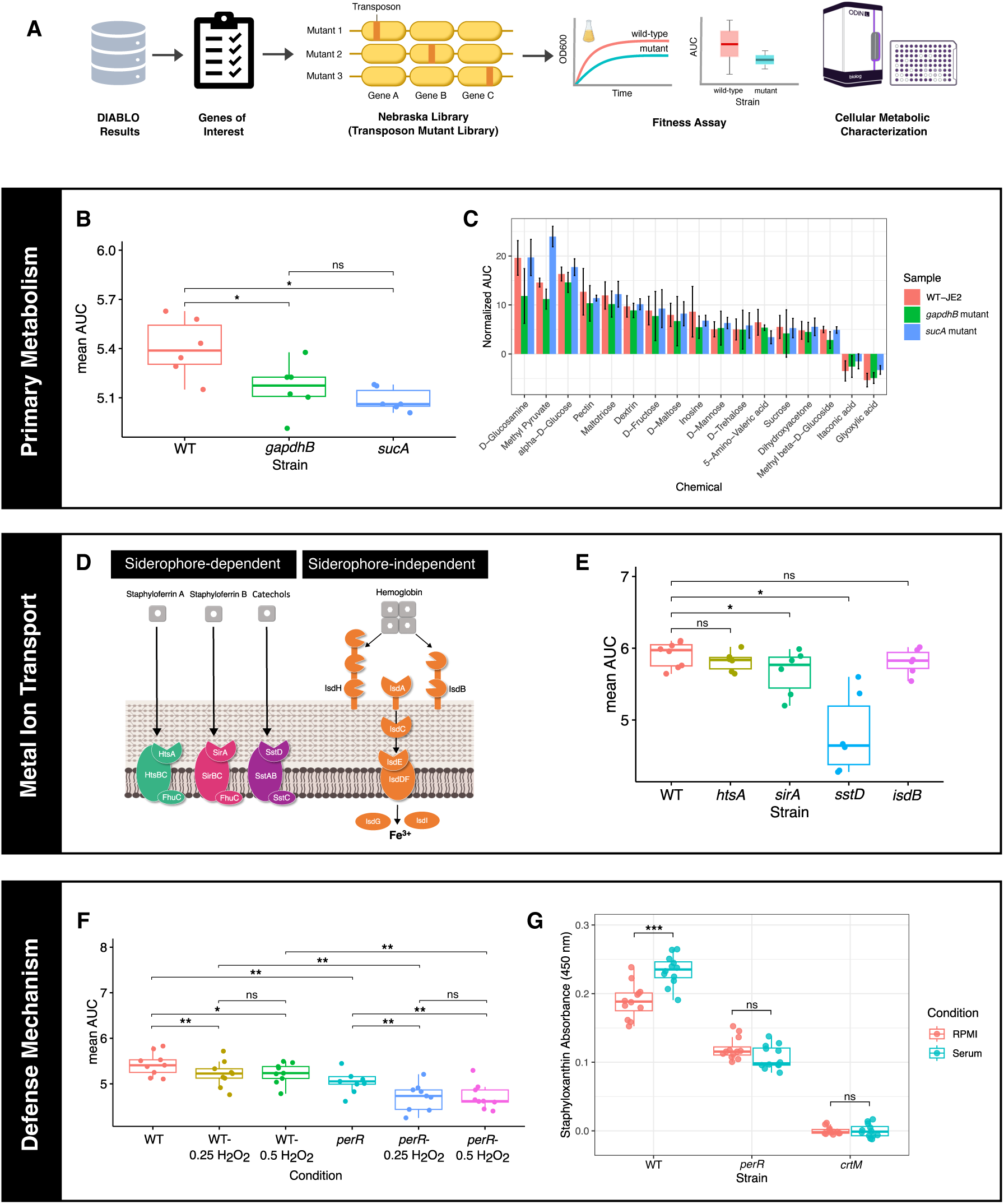
Phenotypic validation of molecular signatures of serum adaptation and metabolic characterization. (A) Workflow of phenotypic validation experiments to confirm the predicted molecular signatures of *S. aureus* exposed to human serum. (B) Box plots comparing Area-under-the-curve (AUC) values of *gapdhB* and *sucA* mutants to wild-type *S. aureus* (WT-JE2) grown in presence of serum, for 20 hours (n = 6 biological replicates). (C) Fitness profiling of *S. aureus* represented by the rate of bacterial respiration in presence of different carbon sources (corresponding to metabolic activity inferred from the reduction of tetrazolium). Plots comparing Area-under-the-curve (AUC) values of *gapdhB* and *sucA* mutants to wild-type *S. aureus* (WT-JE2) grown in presence of sole carbon sources, for 24 hours (n = 3 biological replicates). (D) Representation of iron acquisition pathways in *S. aureus.* (E) Box plots comparing AUC values of iron transport-deficient mutants (*htsA, sirA, sstD, isdB*) to WT-JE2 cultured in serum for 20 hours (n = 6 biological replicates). (F) Box plots comparing AUC values of *perR* mutant to WT-JE2 grown in serum with different concentrations of H_2_O_2_ for 20 hours (n = 9 biological replicates), demonstrating the role of *perR* in oxidative stress resistance. (G) Quantification of Staphyloxanthin production by WT-JE2, *perR* and *crtM* mutants, following 5 h incubation in RPMI or serum (n = 12 replicates, generated from 4 different cultures using 2 serum batches). All data points represent mean values for at least three independent biological replicates, with error bars indicating the corresponding standard deviations from the means. Statistical significance between WT-JE2 and isogenic mutants was determined using the **Wilcoxon paired test**. Significance levels are indicated by asterisks: *p < 0.05, **p < 0.01, and ***p < 0.001.

The DIABLO integrative model identified *gapdhB* and *sucA* as key contributors to response to serum, exhibiting the highest loading values (**Figure 1D, Table S1)**. Disruption of either gene resulted in a significant reduction in bacterial fitness in serum, showing they both play important roles in response to serum exposure (**Figure 3B and S5A**).

GapdhB, a gluconeogenic enzyme, and SucA, an enzyme of the tricarboxylic acid (TCA) cycle, represent distinct nodes in the carbon metabolism pathway. This functional difference suggests that their contributions to *S. aureus* fitness may be contingent upon the availability and type of carbon substrates. To test this, we compared the growth and respiration phenotypes of *gapdhB* and *sucA* transposons mutants with a 190 different carbon sources (Biolog Inc., Hayward, CA, USA), to that of the wild type *S. aureus* JE2 strain. These carbon sources fell into several broad categories, including carbohydrates (mono-, di-, and oligosaccharides), carboxylic acids (short-chain fatty acids like acetate, dicarboxylic acids like fumarate, and hydroxy acids like lactate), amino acids, alcohols, nucleosides and other organic compounds. The metabolic activities of *S. aureus* JE2 wild type (WT), *gapdhB* and *sucA* transposon mutants were more strongly influenced by individual carbon sources, thus compound-specific, than by their broader carbon classification (**Figure S6A**).

Among the tested carbon sources, the carboxylic acid methyl pyruvate, α-D-glucose, D-glucosamine, and other hexoses, significantly enhanced respiration in both WT and mutants, whereas glyoxylic acid and itaconic acid suppressed respiration (**Figure 3C, Figure S6** and **Table S4**). Notably, methyl pyruvate supplementation partially rescued the respiratory deficiency of the *sucA* mutant, surpassing WT levels, while it failed to restore *gapdhB* mutant activity to WT levels (**Figures 3C and S6B**). Whilst we showed that the metabolic responses of WT and mutant strains were shaped more by specific carbon compounds rather than by their general category, supplementation with methyl pyruvate revealed distinct roles for *gapdhB* and *sucA* in pyruvate catabolism (**Figure S6B**). Methyl pyruvate, a membrane permeable pyruvate analog, enter cells where it is converted to acetyl-coA. In the absence of *sucA,* supplementation with methyl pyruvate provides an alternative carbon source that bypasses the glycolytic block caused by the absence of conversion of 2-oxaloglutarate to succinyl-coA in the TCA cycle. GapdhB catalyzes the conversion of glyceraldehyde-3-phosphate (G3P) to 1,3-biphosphoglycerate in the gluconeogenic direction (29). In a *gapdhB* mutant, the gluconeogenic pathway is blocked, so cells cannot convert methyl pyruvate (see primary metabolism in **Figure 4**).

**Figure 4.**
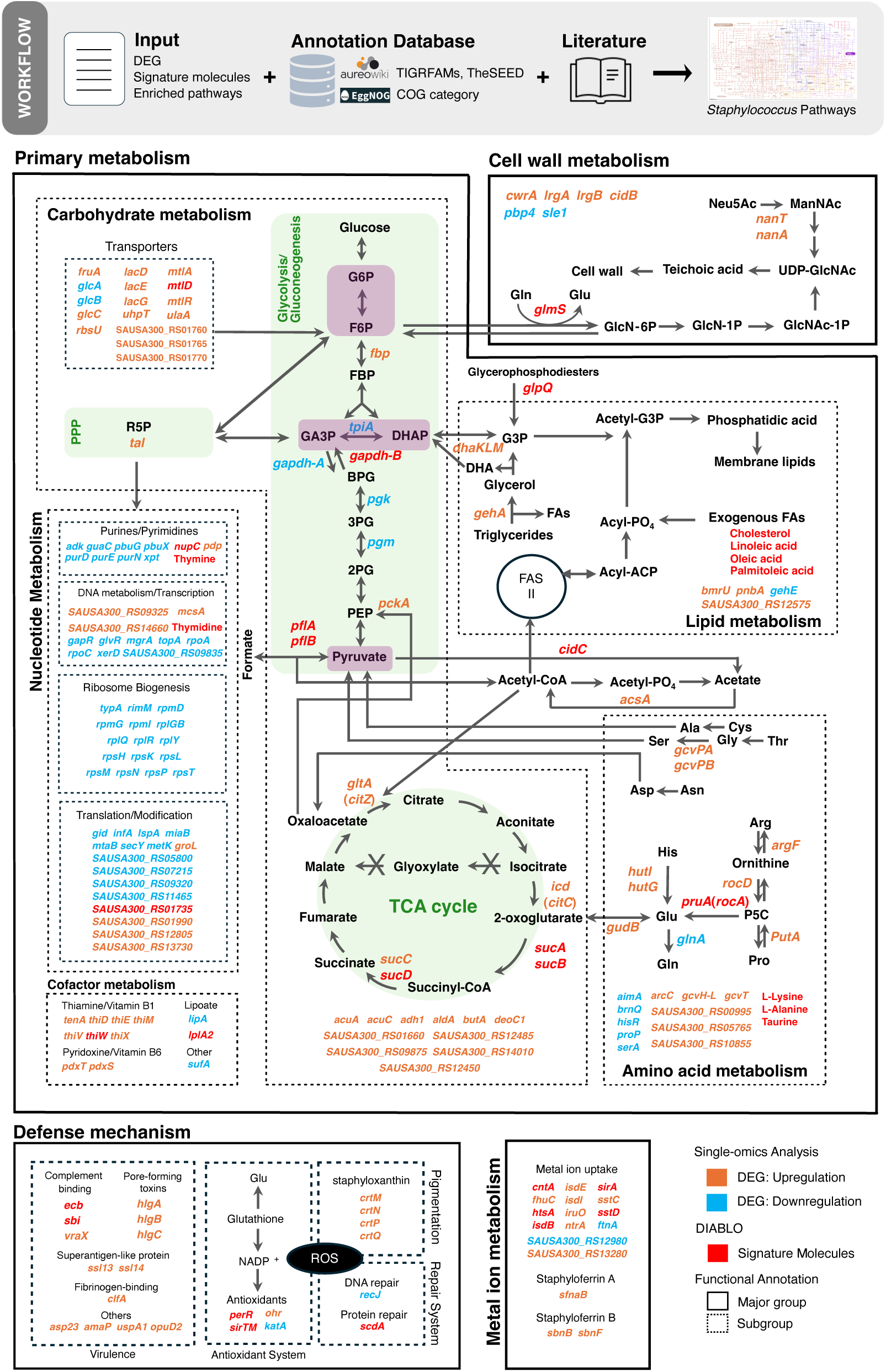
Integrated metabolic and virulence adaptation of *S. aureus* in response to serum. Systems-level metabolic reprogramming of *S. aureus* upon exposure to serum, integrating single- and multi-omics data. The depicted key metabolic pathways include glycolysis/gluconeogenesis (upright rectangle shaded in light green), the pentose phosphate pathway (PPP), the tricarboxylic acid (TCA) cycle (circle shaded in light green), nucleotide biosynthesis, ribosome biogenesis, thiamine (vitamin B1) biosynthesis and transport, cell wall biosynthesis, and lipid metabolism. Host-responsive processes such as immune evasion, metal ion acquisition, and oxidative stress responses are also represented. The name of enzyme is colored to indicate changes in gene expression following serum exposure identified by single omics (compared to RPMI), with orange: increased expression, light blue: decreased expression. Variables predicted by DIABLO (multi-omics analysis) are in red, indicating their status as signature molecules of *S. aureus* serum response.

To investigate the contribution of iron acquisition systems to *Staphylococcus aureus* fitness in serum, we evaluated four mutants, with transposon insertions disrupting genes *htsA*, *sirA*, *sstD*, and *isdB*. The *htsA* and *sirA* genes encode substrate-binding proteins associated with the siderophores staphyloferrin A (SA) and staphyloferrin B (SB), respectively, both of which facilitate iron extraction from transferrin in serum (33). The gene *sstD* encodes a component of the Sst system, which mediates iron uptake from host-derived catechols (33). In contrast, *isdB* encodes a cell wall-anchored hemoglobin receptor within the Isd (iron-regulated surface determinant) system, a siderophore-independent pathway that enables heme iron acquisition (**Figure 3D**). Proteomics revealed significantly elevated levels of HtsA, SirA, SstD, and IsdB following serum exposure, indicating their upregulation in response to iron-limited conditions (**Figure S1C**). *S. aureus sirA* and *sstD* transposon mutants had significantly impaired fitness in serum, consistent with the importance of SB-mediated and catechol-mediated iron uptake under these conditions (**Figure 3E**). In contrast, disruption of *htsA* did not significantly affect fitness, aligned with previous findings that SB biosynthesis, unlike SA, is independent of TCA cycle activity and may be preferentially utilized in iron-restricted environments (55). The reduced fitness observed in the *sirA* mutant likely reflects the impaired utilization of SB as an iron capture process in serum.

Interestingly, the *isdB* mutant exhibited no fitness defect in commercially sourced serum, likely due to the absence of hemoglobin or heme (56), which are typically sequestered within red blood cells and thus unlikely present in commercial cell-free serum preparations. These findings underscore the context-dependent utilization of iron acquisition systems in *S. aureus* and highlight the critical role of siderophore- and catechol-mediated pathways in supporting staphylococcal survival in iron-limited host environments.

Oxidative stress constitutes a fundamental component of the host immune response, primarily through the generation of reactive oxygen species (ROS) aimed at neutralizing invading pathogens. To counteract ROS, bacteria have evolved intricate oxidative stress response mechanisms (57, 58). Central to these systems are redox-sensitive transcriptional regulators such as OhrR, OxyR, and PerR, which enable the detection of specific ROS and orchestrate the staphylococcal defensive gene expression programs (59). In *S. aureus*, PerR functions as a principal peroxide sensor and belongs to the ferric uptake regulator (Fur) family of metalloregulatory proteins, which includes Fur and Zur (60). These regulators require metal cofactors for activity, underscoring the metal-dependent nature of oxidative stress sensing. In our analysis, PerR emerged as a key molecular determinant of oxidative stress resistance in *S. aureus*, a finding confirmed by experimental testing. The *per* mutant exhibited a significant fitness defect in serum supplemented with 0.25- and 0.5 mM hydrogen peroxide (H₂O₂), with the most pronounced impairment observed after 20 hours of exposure to 0.5 mM H₂O₂ (**Figure 3F, Figure S5D**). Also, exposure to serum elicited significant production of staphyloxanthin, encoded by the *crt* operon, and whose production is under the control of PerR (**Figure 3G, Figure S5E**). These results underscore the critical role of PerR in facilitating *S. aureus* survival under oxidative stress conditions, as encountered by *S. aureus* in serum, highlighting its importance in maintaining bacterial fitness in ROS-rich host environments.

### Integrated metabolic responses of *S. aureus* to human serum

To conceptualize the dynamic remodeling of *S. aureus* metabolism in response to serum, we constructed a scaffolded metabolic network based on curated literature (61–74). This network was overlaid with the significant changes in key enzymes identified through single omics analysis (genes upregulated in orange, downregulated in light blue) and by DIABLO integration (genes and metabolites in red), combined with network analyses (**Figure 4, Tables S2–S5**). This integrative approach revealed that *S. aureus* mounts a broad and adaptable metabolic response to the challenges posed by the serum environment, particularly in the utilization of diverse carbon sources including carbohydrates, amino acids, and lipids. Multi-omics data (**Figure S7A–D**) highlighted consistent upregulation of gluconeogenic and TCA cycle enzymes at both transcript and protein levels, indicating a metabolic shift toward energy-efficient carbon utilization upon exposure to serum.

Carbohydrate metabolism was organized around three key metabolic nodes—glucose-6- phosphate/fructose-6-phosphate (G6P/F6P), glyceraldehyde-3-phosphate/dihydroxyacetone phosphate (GA3P/DHAP), and pyruvate (**Figure 4**, purple nodes embedded in glycolysis/gluconeogenesis)—which serve as convergence points for nutrient assimilation. The upregulation of carbohydrate transporter genes (*uhpT, mtlA, fruA, glcC, lacE, SAUSA300_RS01760, rbsU*) in serum aligns with our findings that saccharides, including the hexose monosaccharides α-D-glucose, D-fructose, D-mannose and disaccharides D-maltose, and sucrose, significantly enhanced the metabolic activity of *S. aureus* (**Figures 3C and S6A**). Notably, saccharides such as G6P, fructose, and mannose, but not glucose, have been shown to influence virulence gene expression (75). Our transcriptomic data support this, revealing increased expression of γ-hemolysins genes (*hlgA, hlgB, hlgC*) and immune evasion genes (*ecb, sbi, vraX*) across five *S. aureus* strains in serum. In particular strain BPH2986 (ST8) exhibited strong upregulation of *lukF-PV* and *lukS-PV* genes, encoding Panton-Valentine leucocidins (**Figure S8A**). These findings suggest that carbohydrates, such as fructose and mannose, present in serum, support the metabolic activity of *S. aureus* while increasing virulence gene expression, in contrast to the glucose-rich RPMI.

Amino acids and peptides from serum further contribute to the metabolic adaptability of *S. aureus* by entering carbohydrate metabolism and the TCA cycle through key intermediates such as pyruvate, 2-oxoglutarate, and oxaloacetate. These entry points define three functional groups of amino acids: (1) pyruvate-yielding, (2) 2-oxoglutarate-yielding, and (3) oxaloacetate-yielding (**Figure 4**). Our transcriptomic data revealed significant upregulation of enzymes involved in amino acid catabolism, including *pckA* (encoding pyruvate carboxykinase) and *gudB* (encoding glutamate dehydrogenase), consistent with elevated intracellular levels of phosphoenolpyruvate (PEP) and 2-oxoglutarate (**Figure S7C–E**). Genes encoding enzymes for the degradation of glutamate (*gudB*), proline (*pruA/rocA*, *putA*), arginine (*rocD/argD*), and histidine (*hutI, hutG*) were also upregulated, indicating preferential catabolism of these amino acids during serum exposure. These findings are consistent with previous reports identifying glutamate and its precursors as major carbon sources supporting *S. aureus* proliferation (65), positioning glutamate as a central amino acid in the adaptive response to serum.

Exposure to human serum triggered a marked upregulation of *S. aureus* genes involved in lipid and phospholipid degradation, including *glpQ* and *gehA*. This suggests an enhanced capacity in using host-derived lipids—such as cholesterol, linoleic acid (C18:2), and oleic acid (C18:1)—as nutrient sources. These lipids are metabolized into glycerol-3-phosphate (G3P), which can be then funneled into the carbohydrate metabolism via the GA3P/DHAP node. Interestingly, our transcriptomic data revealed downregulation of *tpiA*, which encodes triosephosphate isomerase, an enzyme responsible for the reversible interconversion of GA3P and DHAP. This suggests a potential bottleneck at this metabolic junction, which may constrain glycolytic flux and influence the efficiency of carbon utilization from lipids present in serum.

Counteracting innate immune defences is essential for *S. aureus* survival during bloodstream infection. Upon exposure to human serum, *S. aureus* exhibited robust transcriptional upregulation of a broad range of virulence-associated genes involved in immune evasion, host colonization, and pathogenesis. These include genes encoding complement-evasion proteins (*ecb, sbi, vraX*), pore-forming cytotoxins (*hlgA, hlgB, hlgC*), superantigen-like proteins (*ssl13, ssl14*), and the fibrinogen-binding adhesin *clfA*. This coordinated immune evasion strategy underscores the pathogen’s capacity to rapidly adapt to immune pressures and enhance its invasive potential in the bloodstream.

Serum exposure also elicited a pronounced oxidative stress response in *S. aureus*. DIABLO analysis identified *perR,* which encodes the peroxide-sensing transcriptional regulator PerR, as a key molecular signature of staphylococcal adaptation to serum. Consistent with this, genes involved in carotenoid biosynthesis —*crtM, crtN, crtQ —*were upregulated, suggesting enhanced anti-oxidative defence through increased production of staphyloxanthin, a carotenoid pigment known to protect against reactive oxygen species (76). In parallel, elevated expression of *scdA*, a gene involved in iron-sulfur cluster repair, indicates activation of protein maintenance pathways essential for preserving enzymatic and cellular functions under oxidative stress (37, 77).

Our integrative approach has highlighted the high adaptive capacity of *S. aureus* to withstand human serum exposure. Through the coordinated activation of nutrient acquisition systems, oxidative stress defenses, and virulence-associated mechanisms, *S. aureus* enhances its ability to survive and persist during systemic infection.

## Discussion

Whilst a leading cause of lethal bacteremia, *S. aureus* must contend with the nutrient-limited and immunologically hostile conditions of the bloodstream to establish infection and cause disease. In the bloodstream, the bacterium uses a large panoply of virulence factors to evade clearance by cellular and humoral immunity (1, 78, 79). Notwithstanding bactericidal immune cells patrolling the bloodstream, serum remains a challenging environment for *S. aureus* due to the paucity of free forms of essential nutrients, often complexed to large proteins (e.g. iron ions complexed to transferrin) (80). Several studies have shown that *S. aureus* has evolved multiple mechanisms, unrelated to its cytotoxicity, to overcome the nutritional constrains and acellular insults of the serum (16). While informing on discrete *S. aureus* nutritional stress responses, these mechanisms were studied in isolation and did not consider the concomitant adaptive responses to the serum environment. Building on our previous work showing that four diverse blood-invasive pathogens, including *S. aureus*, remodeled their cell wall to adapt to serum (18), we used a multi-omics approach to define the shared serum-adaptive response of *S. aureus* across five clinical isolates. Such approach offers a systems-level view of microbial adaptation in response to environmental stimuli, by linking molecular changes across genomic, transcriptomic, proteomic, and metabolomic layers (81–83). Using integrative machine learning-based multivariate analyses, differential expression profiling, and network-based approaches, our study identified staphylococcal reprogramming in cell wall biosynthesis, carbon metabolism, iron transport, and defense mechanisms in response to serum.

The experimental validation of the dominant outputs identified by our integrative framework confirmed the importance of *gapdhB*, *sucA*, *sirA*, *sstD*, and *perR* genes for staphylococcal fitness in serum (**Figure 3**). These results align with the positive co-variation patterns observed in the DIABLO model, suggesting a functional interdependence among the pathways involving these genes (**Figure 1C**). Whilst critical for optimal fitness in serum, *gapdhB* and *sucA* have distinct functions in carbon metabolism. Metabolic profiling revealed that hexose saccharides (e.g., α-D-glucose, D-glucosamine) enhanced the metabolic activity in WT and *gapdhB* and *sucA* mutants, through glycolysis, gluconeogenesis, PPP, and TCA cycle. While methyl pyruvate partially restored bacterial respiration in the *sucA* mutant, likely via conversion to acetyl-CoA and oxaloacetate (84), the *gapdhB* mutant remained metabolically impaired, reflecting incomplete gluconeogenesis and insufficient PPP intermediate generation (**Figures 3B** and **S6**). While highlighting the importance of gluconeogenesis for *S. aureus* during serum exposure, these results further underscore the non-redundant roles of GAPDH homologs (*gapdhA*, *gapdhB*) in maintaining metabolic flux (29). Interestingly, pyruvate was shown to activate *S. aureus* regulatory networks controlling the production of leucocidins, and thus, potentiating the cytotoxicity of *S. aureus* (85). Conversely, exposure to glyoxylate and itaconate inhibited *S. aureus* metabolism **(Figures 3C** and **S6)**. The bacterium does not use the glyoxylate shunt (86), rendering glyoxylate toxic due to metabolic imbalance. Moreover, glyoxylate is limited in serum, as humans lack an active glyoxylate cycle (65). Itaconate, an immunometabolite produced by macrophages, further exacerbates stress by directly inhibiting glycolysis (86, 87). These findings highlight the potential of *S. aureus* metabolic vulnerabilities that could be exploited therapeutically, such as inhibitors of the bacterial glycolysis and the TCA cycle (88). However itaconate was shown to increase tolerance to aminoglycosides, limiting its potential as adjunct therapeutic (89). We constructed a metabolic map of *S. aureus* that integrates the predicted signature molecules elicited/repressed in serum (**Figure 4, Table S4**). This illustrates the flexibility of *S. aureus* in using diverse carbon sources in serum, such as hexose saccharides, amino acids yielding 2-oxoglutarate (e.g., Glu, Pro, Arg, His), and host lipids and fatty acids (e.g., cholesterol, linoleic acid (C18:2), and oleic acid (C18:1)) to support its survival. Hexose saccharides have also been shown to regulate virulence gene expression (75). Peptides and amino acids present in serum contribute to *S. aureus* metabolism by entering glycolysis via pyruvate or intermediates of the TCA cycle (e.g., 2-oxoglutarate, oxaloacetate). Transcriptomic analysis revealed significant upregulation of key enzymes involved in amino acid catabolism, including pyruvate carboxykinase (*pckA*) and glutamate dehydrogenase (*gudB*), phosphoenolpyruvate (PEP) and 2-oxoglutarate (**Figure S7C-E**). Notably, we found a significant increase in gene expression of enzymes (e.g. *gudB*, *pruA* [*rocA*], *putA, rocD [argD], hutI,* and *hutG*) converting amino acids (e.g., Glu, Pro, Arg, His) to 2-oxoglutarate in amino acid catabolism, indicating the usage of these amino acids by the bacteria in serum. This is consistent with the observation that glutamate, and the amino acids that can be converted into glutamate (Pro, Arg, and His), provide most of the carbon required for bacterial growth (65). Several upregulated metabolic pathways facilitating glutamate production, driven by *pruA* (converting proline to glutamate) and *putA* (proline dehydrogenase) underscore glutamate as a key intermediate metabolite in the adaptive response of *S. aureus* to serum. Concurrently, the downregulation of glutamine synthesis, through decreased expression of *glnA*, suggests a reduction in the utilization of glutamate. Excess levels of glutamate did not significantly influence the respiration or growth of JE2 WT (**Table S4**), although several studies have highlighted the important role of glutamate in protecting bacteria from cell death under stress (90–93).

Iron is an important determinant for *S. aureus* fitness (80). Our analyses confirmed the role of multiple iron acquisition mechanisms in iron-limited host environments such as serum. Interestingly, mutants lacking *htsA* and *isdB*, both involved in iron acquisition, displayed growth rates comparable to that of JE2 WT, suggesting some iron uptake system redundancy (**Figure 3E**). This observation is consistent with prior research reporting the upregulation of iron acquisition systems such as Isd and SirABC in *S. aureus* cultured in serum or blood (94). Further studies are necessary to understand the specificity of iron-acquisition modules in iron-limited environments in the context of bacterial metabolic reprograming during bloodstream infection.

Resistance to oxidative stress in serum was also highlighted by our analyses and confirmed experimentally. Staphyloxanthin is a carotenoid pigment produced by *S. aureus* that enhances resistance to oxidative stress. The biosynthesis of staphyloxanthin is encoded by the *crtOPQMN* operon, primarily activated by σB, and its expression is influenced by redox-responsive regulators. PerR, a peroxide-sensing transcriptional repressor identified by DIABLO as a response signature to serum, plays a pivotal role by controlling genes that detoxify reactive oxygen species, such as *katA* that encodes for the catalase. Loss of PerR function disrupts oxidative stress homeostasis, which can alter σB activity, leading to reduced production of staphyloxanthin (**Figure 3G and Figure S5E**). Thus, our data confirmed PerR is a key player linking peroxide sensing to virulence factor regulation upon serum exposure (76).

Serum exposure also engaged *S. aureus* defense mechanisms against acellular immunity. We observed upregulation of virulence-related genes including complement-binding proteins (e.g., *ecb*, *sbi*, *vraX*) (28, 95, 96), pore-forming toxins (e.g., the γ-hemolysins *hlgA*, *hlgB*, *hlgC*) (97), superantigen-like proteins (*ssl13*, *ssl14*) (98), and the fibrinogen-binding protein *clfA* (99) (**Figures 4 and S8)**. Such responses would prime *S. aureus* against clearance by immune cells. Mechanisms for evasion from host antibodies and the complement system were also engaged, including the immunoglobulin-binding protein (Sbi) and complement convertase inhibitor (Ecb) (28, 100). *S. aureus* sensing and responses to host oxidative stress were also identified as important for serum exposure survival (40, 43, 47, 60). Future studies examining how serum promotes *S. aureus* biofilm formation on host-like surfaces could provide deeper insight into the roles of ClfA, Ecb and other adherence factors during bloodstream infections.

### Conclusion and perspectives

This study exploited multi-omics analyses, including machine learning-based approaches, DE analysis and network analysis, to uncover the major *S. aureus* features engaged in response to human serum. Our findings highlight the parallel activation of nutrient acquisition systems, oxidative stress defenses, and virulence-associated mechanisms, underlining the strong adaptive capacity of *S. aureus* to withstand the multifactorial challenges imposed by human serum. While serum is acellular and does not reflect the breadth of host-pathogen interactions during bloodstream infections, serum can be an informative proxy for understanding how invading bacteria survive in blood. The key staphylococcal pathways identified here could be further dissected, to explore staphylococcal adaptive responses to identify prospective bacterial (and host) targets for new therapeutic approaches.

### Limitations of the study

The integrative analysis performed here was highly sensitive and detected subtle patterns linking different omic datasets. However, the metabolic flexibility and functional redundancy within key bacterial pathways - particularly central carbon metabolism and iron acquisition – are likely not limited to individual contributions of specific genes, thus making genes contributing to these pathways difficult to confirm experimentally using single isogenic mutants. Furthermore, the individual contributions of metabolites such as citrate, an intermediate of the TCA cycle, can repress the activity of the transcriptional activity of genes involved in iron homeostasis (101), underscoring the cross-talks between pathways that could alter single -omic readouts, and thus multi-omics analyses. This study used commercially sourced serum for phenotypic confirmation of the staphylococcal signatures. Whilst such reagent offers more control, such as batch control, the concentrations of carbohydrates, metal ions, and reactive oxygen species are unknown, which may influence the significance of phenotypic outputs. Moreover, the datasets analyzed here were generated using pooled human sera (Lifeblood, Melbourne, Australia), whilst our phenotypic confirmation experiments were conducted using commercially sourced and batch-controlled human sera (Sigma Aldrich). The difference in serum processing may explain the discrepant phenotypes obtained for the siderophore-independent *isd* genes involved in iron-scavenging from heme between our current study and the study that generated the datasets (18). Whilst RPMI, closely mimicking the nutrient-limited of the milieu encountered by *S. aureus* during host infection (102), was used as the comparator to serum in our studies ((18) and present study), bacterial omics analyses generated from exposure to alternative media, such as the Human Like Plasma Media (HPLM) (103), enriched in host-like metabolites, could refine the staphylococcal signatures in future analyses.

## METHODS

### *S. aureus* genome and pangenome analyses

The complete genomes of five clinical *S. aureus* strains, BPH2760, BPH2819, BPH2900, BPH2947, BPH2986 (18), and of the strain USA300 JE2 were downloaded from the NCBI database (https://www.ncbi.nlm.nih.gov/bioproject/PRJEB29881/, https://www.ncbi.nlm.nih.gov/bioproject/PRJNA381486).

Genomic analysis was conducted using Bohra v2.3.6 (https://github.com/MDU-PHL/bohra). Bohra provided information such as Single Nucleotide Polymorphisms (SNPs), phylogeny, multi-locus sequence typing (MLST), resistome, virulome, and Pan-genome. We used Phandango to visualize the pangenome profiles (https://jameshadfield.github.io/phandango/#/).

The AureoWiki database (24) was used to annotate *S. aureus* genes/proteins and to define orthologous relationships, through a reciprocal best hit (RBH) search using MMseqs2 with default parameters (https://github.com/soedinglab/MMseqs2) between the five studied genomes (FAA format) and the *S. aureus* reference genome and proteome (USA300_FPR3757_NCBI_2017 and USA300_FPR3757_NCBI_UniProt_2013).

### Confirmation of *S. aureus* transposon mutants’ genome sequence

We used *S. aureus* transposon mutants of signature genes identified by our analytical pipeline, to verify their fitness in serum. These mutants were selected from the Nebraska transposon library (NTL), including NE275, NE400, NE547, NE665, NE1102, NE1343, and NE1767 (104). The mutants are expected to contain a precise and non-polar transposon insertions, without additional mutations. All mutants were genome sequenced on the Oxford Nanopore platform using Min114 flow cells. Base calling, demultiplexing, and adapter removal were performed in real-time using MinKNOW. Mutant NE1102 was sequenced on a GridION and processed using Guppy. Remaining mutants were sequenced on a MinION and processed using Dorado. FASTQ reads were assembled using Trycycler version 0.5.3 (105). The assembled genomes were compared to the cognate wild-type reference genome using the Mauve whole-genome aligner plugin tool in the Geneious Prime (version 2022.2.2) (106).

### Serum treatment

We used batch-controlled human serum (Male AB plasma, Sigma Aldrich) to reduce variations arising from different donors. Human serum was clarified from lipids by centrifugation at 3,000 x g during 5 min at 4°C. We further analyzed the protein content of clarified serum (**Figure S9**). Human serum samples (pre (untreated, U) and post (treated, T) centrifugation 5 min at 3000 x g) were diluted 1:50 in sterile PBS, then boiled 5 minutes with 5mM DTT in 1x protein loading buffer. Equivalent amounts of proteins were loaded on a Bolt 4%-12% Bis-Tris SDS PAGE gel and separated using with MOPS-SDS running buffer (ThermoFisher, USA) at 100 volts for 1h. The gel was stained with Coomassie blue overnight, destained and imaged using a GE Amersham Imager 800 (GE Life Sciences, UK).

### *S. aureus* fitness in serum

Overnight cultures of *S. aureus* JE2 (WT), and NTL mutants of *gapdhB*, *sucA*, *htsA*, *sirA*, *sstD*, *isdB* and *perR* were grown from single colonies in 10 ml Brain Heart Infusion media (BHI, BD BACTO^TM^) at 37 °C with orbital shaking. Overnight cultures were centrifuged for 5 min at 5,000 x g, and pellets were washed twice with sterile PBS. The cells were then standardized to an OD 600 nm = 0.5 (0.5 UOD) in RPMI (Gibco^TM^). Bacteria equivalent to 0.01 UOD were inoculated in 180 μL of BHI, RPMI or 50% serum (diluted in RPMI) in a clear 96-well plate (Costar), in triplicate to a final volume of 200 μL. Plates were incubated at 37°C in ClariostarPLUS microplate reader (BMG Labtech) and bacterial growth was measured every 10 min for 20 h at 600 nm.

### Phenotypic analysis of *S. aureus* carbohydrate metabolism

Phenotypic profiling was performed using the Biolog Phenotype MicroArray™ (PM) system with PM1 and PM2 plates (Biolog Inc., Hayward, CA, USA), to determine the *S. aureus* fitness in presence of 190 distinct carbon sources. Overnight *S. aureus* cultures were grown in BHI at 37 °C with shaking. Bacterial suspensions were prepared according to the manufacturer’s protocol for Gram-positive organisms. The inocula of *gapdhB*, *sucA* and parental WT strain were loaded into PM1 and PM2 plates and incubated at 37 °C for 24 hours in the Biolog Odin™ system, which enables the simultaneous monitoring of growth and respiration (by redox dye reduction). Metabolic activity was assessed by measuring the reduction of tetrazolium-based redox dyes, which serve as indicators of electron transport chain activity in response to specific carbon sources. Dye reduction results in the formation of purple formazan, with absorbance recorded every 20 minutes at 590 nm. The area under the curve (AUC) for each well was calculated to quantify metabolic activity, enabling comparative analysis of carbon source utilization across conditions or strains.

### Quantification of Staphyloxanthin

Overnight cultures of *S. aureus* JE2 (WT), and NTL mutants of *perR* and *crtM* were grown from single colonies in 25 ml BHI, at 37 °C with orbital shaking. The cultures were standardized to an OD 600 nm = 25. These were then centrifuged for 15 min at 5,000 x g, and pellets were washed twice with sterile PBS. Bacteria were incubated in 700 μL of RPMI or in 50% serum (diluted in RPMI) at 37 °C with orbital shaking for 5 h. The bacteria were then centrifuged for 5 min at 5,000 x g, and the supernatant was discarded. The pellets were resuspended in 450 μL of methanol, transferred in 1.5 ml tubes and incubated at 55 °C for 15 min with shaking. Tubes were centrifuged for 2 min at 13,000 RPM, at room temperature. The 100 μL of supernatant from each tube was added to a clear 96-well plate (Costar) in triplicate and extracted staphyloxanthin was quantified at 450 nm using a ClariostarPLUS microplate reader (BMG Labtech). Staphyloxanthin production was calculated by subtracting each data point from the mean absorbance of *crtM* to normalize different condition backgrounds (RPMI and Serum). Replicates were generated from 4 different cultures for each strain, assessed in technical triplicate across two distinct commercial serum batches.

### Principal component analysis

Principal Component Analysis (PCA) was independently performed on each omics dataset— transcriptomics, proteomics, GC-MS metabolomics, and LC-MS metabolomics—to explore global data structure and identify the primary sources of variation between serum- and RPMI-exposed samples. PCA was conducted using the *pca()* function from the mixOmics package (v6.24.0) in R (v4.2.3), with data scaled to unit variance. The optimal number of components was determined using the tune.pca() function, which indicated that the first three components captured approximately 50% of the total variance across datasets.

### Multi-omics data integration

We implemented the MixOmics data analytic pipeline known as Multiblock (s)PLS-DA or DIABLO, a multivariate statistical method for classifying and separating different categorical classes (19, 20). This supervised multi-omics integrative method adds sparsity to the model, improving result interpretability by selecting the most relevant variables and setting others’ coefficients to zero. DIABLO treats each omics dataset as a separate “block” and identifies co-variation patterns across these blocks, to determine interactions between, and changes by, biological molecules under various conditions. The DIABLO model was designed with a selected design matrix weight of 0.5 to balance discrimination between conditions and correlation across omics layers, ensuring that feature selection prioritized both inter-omics agreement and RPMI-Serum separation. An initial model trained with five components revealed that the first component alone was sufficient to clearly discriminate between serum and RPMI conditions. To assess model robustness and guide component selection, we performed leave-one-out (LOO) cross-validation, which confirmed that a single component provided perfect classification performance. To optimize feature selection for each omics layer, we performed parameter tuning using the tune.block.splsda function. A grid of candidate values for the number of features to retain per block was defined: 20–100 (step size 20) for transcriptomics and proteomics, and 10–100 (step size 10) for both metabolomics datasets (GC-MS and LC-MS). The model was tuned using 5-fold cross-validation repeated 10 times and centroid distance as the classification metric. The number of features retained per block was: 20 for transcriptomics, 20 for proteomics, 10 for GC-MS, and 10 for LC-MS, totaling 60 features. Using these parameters, the final DIABLO model was built using block.splsda function with one component (as determined during component tuning)(19, 20).

### Enrichment analysis

To identify staphylococcal pathways prominently represented following exposure to serum, feature lists derived from all omics data underwent pathway enrichment analysis, using over-representation analysis (ORA) and Gene Set Enrichment Analysis (GSEA). Pathway enrichment analysis was performed with package clusterProfiler v4.6.2. The obtained feature lists were mapped to the associated biological annotation terms (e.g. GO terms or KEGG pathways) and statistical tests for enrichment probability (enrichment P-value) were calculated to obtain the enrichment of the feature sets. In ORA, the preselected features were filtered by a criterion of log2FC > 1 and p-value < 0.05. For GSEA, all features were ranked based on their log2FC values, meaning features with the largest positive and negative expression changes are positioned at opposite ends of the ranked list, while those with smaller or no changes are located near the middle.

### Network analysis

The biological relationships among the list of enriched pathways were identified by selecting the significant pathways with p.adjust < 0.05 (except for metabolomic datasets; p-value < 0.05). Further network analysis, using pairwise_termsim function in the enrichplot package in R, calculated the pairwise similarity of the enriched terms using Jaccard’s similarity coefficient or the similarity of gene subsets sharing between pathways. Conversion to enrichment map to dataframes provided similarity scores and number of strains sharing individual enriched pathways, represented as node and edge, respectively, for networks’ visualization.

### Data availability

All codes are available on GitHub (https://github.com/warasinee/Multiomics_Analyses_2024). Oxford Nanopore reads of all strains and mutants generated are deposited in NCBI SRA (BioProject ID PRJNA1284587).

## Acknowledgements

This research was supported by the Development and Promotion of Science and Technology Talents Project (DPST) from the Thai Government (WM), the National Health and Medical Research Council of Australia to BPH (GNT1196103), TPS (GNT1194325), AH, SGG, RG (GNT2018880).

## References

1. Howden BP, Giulieri SG, Wong Fok Lung T, Baines SL, Sharkey LK, Lee JYH, et al. Staphylococcus aureus host interactions and adaptation. Nat Rev Microbiol. 2023;21(6):380–95.

2. Räz AK, Andreoni F, Boumasmoud M, Bergada-Pijuan J, Schweizer TA, Mairpady Shambat S, et al. Limited Adaptation of Staphylococcus aureus during Transition from Colonization to Invasive Infection. Microbiol Spectr. 2023;11(4):e0259021.

3. Murray CJ, Ikuta KS, Sharara F, Swetschinski L, Aguilar GR, Gray A, et al. Global burden of bacterial antimicrobial resistance in 2019: a systematic analysis. The lancet. 2022;399(10325):629–55.

4. Guerra FE, Borgogna TR, Patel DM, Sward EW, Voyich JM. Epic Immune Battles of History: Neutrophils vs. Staphylococcus aureus. Front Cell Infect Microbiol. 2017;7:286.

5. Hornef MW, Wick MJ, Rhen M, Normark S. Bacterial strategies for overcoming host innate and adaptive immune responses. Nat Immunol. 2002;3(11):1033–40.

6. Mathew J, Sankar P, Varacallo M. Physiology, Blood Plasma. StatPearls. Treasure Island (FL): StatPearls Publishing Copyright © 2023, StatPearls Publishing LLC.; 2023.

7. Kuehl R, Morata L, Meylan S, Mensa J, Soriano A. When antibiotics fail: a clinical and microbiological perspective on antibiotic tolerance and persistence of Staphylococcus aureus. J Antimicrob Chemother. 2020;75(5):1071–86.

8. Lopatkin AJ, Stokes JM, Zheng EJ, Yang JH, Takahashi MK, You L, et al. Bacterial metabolic state more accurately predicts antibiotic lethality than growth rate. Nat Microbiol. 2019;4(12):2109–17.

9. Price EE, Boyd JM. Genetic Regulation of Metal Ion Homeostasis in Staphylococcus aureus. Trends Microbiol. 2020;28(10):821–31.

10. Kwiecinski JM, Horswill AR. Staphylococcus aureus bloodstream infections: pathogenesis and regulatory mechanisms. Curr Opin Microbiol. 2020;53:51–60.

11. Bear A, Locke T, Rowland-Jones S, Pecetta S, Bagnoli F, Darton TC. The immune evasion roles of Staphylococcus aureus protein A and impact on vaccine development. Front Cell Infect Microbiol. 2023;13:1242702.

12. de Jong NWM, van Kessel KPM, van Strijp JAG. Immune Evasion by Staphylococcus aureus. Microbiol Spectr. 2019;7(2).

13. Paiva TO, Geoghegan JA, Dufrêne YF. High-force catch bonds between the Staphylococcus aureus surface protein SdrE and complement regulator factor H drive immune evasion. Commun Biol. 2023;6(1):302.

14. Joo HS, Otto M. Mechanisms of resistance to antimicrobial peptides in staphylococci. Biochim Biophys Acta. 2015;1848(11 Pt B):3055–61.

15. Ledger EVK, Mesnage S, Edwards AM. Human serum triggers antibiotic tolerance in Staphylococcus aureus. Nat Commun. 2022;13(1):2041.

16. Murdoch CC, Skaar EP. Nutritional immunity: the battle for nutrient metals at the host-pathogen interface. Nat Rev Microbiol. 2022;20(11):657–70.

17. Cassat JE, Skaar EP. Metal ion acquisition in Staphylococcus aureus: overcoming nutritional immunity. Semin Immunopathol. 2012;34(2):215–35.

18. Mu A, Klare WP, Baines SL, Ignatius Pang CN, Guérillot R, Harbison-Price N, et al. Integrative omics identifies conserved and pathogen-specific responses of sepsis-causing bacteria. Nat Commun. 2023;14(1):1530.

19. Rohart F, Gautier B, Singh A, KA LC. mixOmics: An R package for ’omics feature selection and multiple data integration. PLoS Comput Biol. 2017;13(11):e1005752.

20. Singh A, Shannon CP, Gautier B, Rohart F, Vacher M, Tebbutt SJ, et al. DIABLO: an integrative approach for identifying key molecular drivers from multi-omics assays. Bioinformatics. 2019;35(17):3055–62.

21. González I, Cao KA, Davis MJ, Déjean S. Visualising associations between paired ’omics’ data sets. BioData Min. 2012;5(1):19.

22. Cox M, Nelson D. Lehninger Principles of Biochemistry. 5 ed: W. H. Freeman and Company; 2008.

23. Simon GM, Cravatt BF. Activity-based proteomics of enzyme superfamilies: serine hydrolases as a case study. J Biol Chem. 2010;285(15):11051–5.

24. Fuchs S, Mehlan H, Bernhardt J, Hennig A, Michalik S, Surmann K, et al. AureoWiki ̵ The repository of the Staphylococcus aureus research and annotation community. Int J Med Microbiol. 2018;308(6):558–68.

25. Ghssein G, Brutesco C, Ouerdane L, Fojcik C, Izaute A, Wang S, et al. Biosynthesis of a broad-spectrum nicotianamine-like metallophore in Staphylococcus aureus. Science. 2016;352(6289):1105–9.

26. Grim KP, San Francisco B, Radin JN, Brazel EB, Kelliher JL, Párraga Solórzano PK, et al. The Metallophore Staphylopine Enables Staphylococcus aureus To Compete with the Host for Zinc and Overcome Nutritional Immunity. mBio. 2017;8(5).

27. Remy L, Carrière M, Derré-Bobillot A, Martini C, Sanguinetti M, Borezée-Durant E. The Staphylococcus aureus Opp1 ABC transporter imports nickel and cobalt in zinc-depleted conditions and contributes to virulence. Mol Microbiol. 2013;87(4):730–43.

28. Jongerius I, Garcia BL, Geisbrecht BV, van Strijp JAG, Rooijakkers SHM. Convertase inhibitory properties of Staphylococcal extracellular complement-binding protein. J Biol Chem. 2010;285(20):14973–9.

29. Purves J, Cockayne A, Moody PC, Morrissey JA. Comparison of the regulation, metabolic functions, and roles in virulence of the glyceraldehyde-3-phosphate dehydrogenase homologues gapA and gapB in Staphylococcus aureus. Infect Immun. 2010;78(12):5223–32.

30. Jorge AM, Schneider J, Unsleber S, Göhring N, Mayer C, Peschel A. Utilization of glycerophosphodiesters by Staphylococcus aureus. Mol Microbiol. 2017;103(2):229–41.

31. Gustafson J, Strässle A, Hächler H, Kayser FH, Berger-Bächi B. The femC locus of Staphylococcus aureus required for methicillin resistance includes the glutamine synthetase operon. J Bacteriol. 1994;176(5):1460–7.

32. Wolff DW, Deng Z, Bianchi-Smiraglia A, Foley CE, Han Z, Wang X, et al. Phosphorylation of guanosine monophosphate reductase triggers a GTP-dependent switch from pro- to anti-oncogenic function of EPHA4. Cell Chem Biol. 2022;29(6):970–84.e6.

33. Beasley FC, Marolda CL, Cheung J, Buac S, Heinrichs DE. Staphylococcus aureus transporters Hts, Sir, and Sst capture iron liberated from human transferrin by Staphyloferrin A, Staphyloferrin B, and catecholamine stress hormones, respectively, and contribute to virulence. Infect Immun. 2011;79(6):2345–55.

34. Mathelié-Guinlet M, Viela F, Alfeo MJ, Pietrocola G, Speziale P, Dufrêne YF. Single-Molecule Analysis Demonstrates Stress-Enhanced Binding between Staphylococcus aureus Surface Protein IsdB and Host Cell Integrins. Nano Lett. 2020;20(12):8919–25.

35. Alfeo MJ, Pagotto A, Barbieri G, Foster TJ, Vanhoorelbeke K, De Filippis V, et al. Staphylococcus aureus iron-regulated surface determinant B (IsdB) protein interacts with von Willebrand factor and promotes adherence to endothelial cells. Sci Rep. 2021;11(1):22799.

36. Burman JD, Leung E, Atkins KL, O’Seaghdha MN, Lango L, Bernadó P, et al. Interaction of human complement with Sbi, a staphylococcal immunoglobulin-binding protein: indications of a novel mechanism of complement evasion by Staphylococcus aureus. J Biol Chem. 2008;283(25):17579–93.

37. Overton TW, Justino MC, Li Y, Baptista JM, Melo AM, Cole JA, et al. Widespread distribution in pathogenic bacteria of di-iron proteins that repair oxidative and nitrosative damage to iron-sulfur centers. J Bacteriol. 2008;190(6):2004–13.

38. Brunskill EW, de Jonge BLM, Bayles KW. The Staphylococcus aureus scdA gene: a novel locus that affects cell division and morphogenesis. Microbiology (Reading). 1997;143 (Pt 9):2877–82.

39. Zhang W, Zhang M, Gao C, Zhang Y, Ge Y, Guo S, et al. Coupling between d-3-phosphoglycerate dehydrogenase and d-2-hydroxyglutarate dehydrogenase drives bacterial l-serine synthesis. Proc Natl Acad Sci U S A. 2017;114(36):E7574–e82.

40. Rack JG, Morra R, Barkauskaite E, Kraehenbuehl R, Ariza A, Qu Y, et al. Identification of a Class of Protein ADP-Ribosylating Sirtuins in Microbial Pathogens. Mol Cell. 2015;59(2):309–20.

41. Arragain S, Handelman SK, Forouhar F, Wei FY, Tomizawa K, Hunt JF, et al. Identification of eukaryotic and prokaryotic methylthiotransferase for biosynthesis of 2-methylthio-N6-threonylcarbamoyladenosine in tRNA. J Biol Chem. 2010;285(37):28425–33.

42. Murphey WH, Rosenblum ED. MANNITOL CATABOLISM BY STAPHYLOCOCCUS AUREUS. Arch Biochem Biophys. 1964;107:292–7.

43. Ji CJ, Kim JH, Won YB, Lee YE, Choi TW, Ju SY, et al. Staphylococcus aureus PerR Is a Hypersensitive Hydrogen Peroxide Sensor using Iron-mediated Histidine Oxidation. J Biol Chem. 2015;290(33):20374–86.

44. Leibig M, Liebeke M, Mader D, Lalk M, Peschel A, Götz F. Pyruvate formate lyase acts as a formate supplier for metabolic processes during anaerobiosis in Staphylococcus aureus. J Bacteriol. 2011;193(4):952–62.

45. Bertrand BP, Heim CE, West SC, Chaudhari SS, Ali H, Thomas VC, et al. Role of Staphylococcus aureus Formate Metabolism during Prosthetic Joint Infection. Infect Immun. 2022;90(11):e0042822.

46. Handa N, Terada T, Doi-Katayama Y, Hirota H, Tame JR, Park SY, et al. Crystal structure of a novel polyisoprenoid-binding protein from Thermus thermophilus HB8. Protein Sci. 2005;14(4):1004–10.

47. Rack JGM, Palazzo L, Ahel I. (ADP-ribosyl)hydrolases: structure, function, and biology. Genes Dev. 2020;34(5-6):263–84.

48. Christgen SL, Becker DF. Role of Proline in Pathogen and Host Interactions. Antioxid Redox Signal. 2019;30(4):683–709.

49. Richardson AR, Somerville GA, Sonenshein AL. Regulating the Intersection of Metabolism and Pathogenesis in Gram-positive Bacteria. Microbiol Spectr. 2015;3(3).

50. Kim J, Kim GL, Norambuena J, Boyd JM, Parker D. Impact of the pentose phosphate pathway on metabolism and pathogenesis of Staphylococcus aureus. PLoS Pathog. 2023;19(7):e1011531.

51. Njenga R, Boele J, Öztürk Y, Koch HG. Coping with stress: How bacteria fine-tune protein synthesis and protein transport. J Biol Chem. 2023;299(9):105163.

52. Gibbs MR, Moon KM, Warner BR, Chen M, Bundschuh R, Foster LJ, et al. Functional Analysis of BipA in E. coli Reveals the Natural Plasticity of 50S Subunit Assembly. J Mol Biol. 2020;432(19):5259–72.

53. Esquilin-Lebron K, Dubrac S, Barras F, Boyd JM. Bacterial Approaches for Assembling Iron-Sulfur Proteins. mBio. 2021;12(6):e0242521.

54. Barrault M, Leclair E, Kumeko EK, Jacquet E, Bouloc P. Staphylococcal sRNA IsrR downregulates methylthiotransferase MiaB under iron-deficient conditions. Microbiol Spectr. 2024;12(10):e0388823.

55. Sheldon JR, Marolda CL, Heinrichs DE. TCA cycle activity in Staphylococcus aureus is essential for iron-regulated synthesis of staphyloferrin A, but not staphyloferrin B: the benefit of a second citrate synthase. Mol Microbiol. 2014;92(4):824–39.

56. Pishchany G, Sheldon JR, Dickson CF, Alam MT, Read TD, Gell DA, et al. IsdB-dependent hemoglobin binding is required for acquisition of heme by Staphylococcus aureus. J Infect Dis. 2014;209(11):1764–72.

57. Imlay JA. Cellular defenses against superoxide and hydrogen peroxide. Annu Rev Biochem. 2008;77:755–76.

58. Toledano MB, Delaunay A, Monceau L, Tacnet F. Microbial H2O2 sensors as archetypical redox signaling modules. Trends Biochem Sci. 2004;29(7):351–7.

59. Dubbs JM, Mongkolsuk S. Peroxide-sensing transcriptional regulators in bacteria. J Bacteriol. 2012;194(20):5495–503.

60. Horsburgh MJ, Clements MO, Crossley H, Ingham E, Foster SJ. PerR controls oxidative stress resistance and iron storage proteins and is required for virulence in Staphylococcus aureus. Infect Immun. 2001;69(6):3744–54.

61. Hines KM, Alvarado G, Chen X, Gatto C, Pokorny A, Alonzo F, 3rd, et al. Lipidomic and Ultrastructural Characterization of the Cell Envelope of Staphylococcus aureus Grown in the Presence of Human Serum. mSphere. 2020;5(3).

62. Beasley FC, Cheung J, Heinrichs DE. Mutation of L-2,3-diaminopropionic acid synthase genes blocks staphyloferrin B synthesis in Staphylococcus aureus. BMC Microbiol. 2011;11:199.

63. Carvalho SM, de Jong A, Kloosterman TG, Kuipers OP, Saraiva LM. The Staphylococcus aureus α-Acetolactate Synthase ALS Confers Resistance to Nitrosative Stress. Front Microbiol. 2017;8:1273.

64. Gaupp R, Ledala N, Somerville GA. Staphylococcal response to oxidative stress. Front Cell Infect Microbiol. 2012;2:33.

65. Halsey CR, Lei S, Wax JK, Lehman MK, Nuxoll AS, Steinke L, et al. Amino Acid Catabolism in Staphylococcus aureus and the Function of Carbon Catabolite Repression. mBio. 2017;8(1).

66. Imada A, Nozaki Y, Kawashima F, Yoneda M. Regulation of glucosamine utilization in Staphylococcus aureus and Escherichia coli. J Gen Microbiol. 1977;100(2):329–37.

67. Kenny JG, Moran J, Kolar SL, Ulanov A, Li Z, Shaw LN, et al. Mannitol utilisation is required for protection of Staphylococcus aureus from human skin antimicrobial fatty acids. PLoS One. 2013;8(7):e67698.

68. Kriegeskorte A, Block D, Drescher M, Windmüller N, Mellmann A, Baum C, et al. Inactivation of thyA in Staphylococcus aureus attenuates virulence and has a strong impact on metabolism and virulence gene expression. mBio. 2014;5(4):e01447–14.

69. Nolan AC, Zeden MS, Kviatkovski I, Campbell C, Urwin L, Corrigan RM, et al. Purine Nucleosides Interfere with c-di-AMP Levels and Act as Adjuvants To Re-Sensitize MRSA To β-Lactam Antibiotics. mBio. 2023;14(1):e0247822.

70. O’Connor MJ, Bartler AV, Ho KC, Zhang K, Casas Fuentes RJ, Melnick BA, et al. Understanding Staphylococcus aureus in hyperglycaemia: A review of virulence factor and metabolic adaptations. Wound Repair Regen. 2024;32(5):661–70.

71. Parsons JB, Rock CO. Bacterial lipids: metabolism and membrane homeostasis. Prog Lipid Res. 2013;52(3):249–76.

72. Potter AD, Butrico CE, Ford CA, Curry JM, Trenary IA, Tummarakota SS, et al. Host nutrient milieu drives an essential role for aspartate biosynthesis during invasive Staphylococcus aureus infection. Proc Natl Acad Sci U S A. 2020;117(22):12394–401.

73. Qi Z, Sun N, Liu C. Glyoxylate cycle maintains the metabolic homeostasis of Pseudomonas aeruginosa in viable but nonculturable state induced by chlorine stress. Microbiol Res. 2023;270:127341.

74. Sachla AJ, Helmann JD. A bacterial checkpoint protein for ribosome assembly moonlights as an essential metabolite-proofreading enzyme. Nat Commun. 2019;10(1):1526.

75. Seo KS, Park N, Rutter JK, Park Y, Baker CL, Thornton JA, et al. Role of Glucose-6-Phosphate in Metabolic Adaptation of Staphylococcus aureus in Diabetes. Microbiol Spectr. 2021;9(2):e0085721.

76. Clauditz A, Resch A, Wieland KP, Peschel A, Götz F. Staphyloxanthin plays a role in the fitness of Staphylococcus aureus and its ability to cope with oxidative stress. Infect Immun. 2006;74(8):4950–3.

77. Chang W, Small DA, Toghrol F, Bentley WE. Global transcriptome analysis of Staphylococcus aureus response to hydrogen peroxide. J Bacteriol. 2006;188(4):1648–59.

78. Cheung GYC, Bae JS, Otto M. Pathogenicity and virulence of Staphylococcus aureus. Virulence. 2021;12(1):547–69.

79. Soe YM, Bedoui S, Stinear TP, Hachani A. Intracellular Staphylococcus aureus and host cell death pathways. Cell Microbiol. 2021;23(5):e13317.

80. Ledala N, Zhang B, Seravalli J, Powers R, Somerville GA. Influence of iron and aeration on Staphylococcus aureus growth, metabolism, and transcription. J Bacteriol. 2014;196(12):2178–89.

81. Fouché A, Zinovyev A. Omics data integration in computational biology viewed through the prism of machine learning paradigms. Front Bioinform. 2023;3:1191961.

82. Maghsoudi Z, Nguyen H, Tavakkoli A, Nguyen T. A comprehensive survey of the approaches for pathway analysis using multi-omics data integration. Brief Bioinform. 2022;23(6).

83. Vahabi N, Michailidis G. Unsupervised Multi-Omics Data Integration Methods: A Comprehensive Review. Front Genet. 2022;13:854752.

84. Campbell C, Fingleton C, Zeden MS, Bueno E, Gallagher LA, Shinde D, et al. Accumulation of Succinyl Coenzyme A Perturbs the Methicillin-Resistant Staphylococcus aureus (MRSA) Succinylome and Is Associated with Increased Susceptibility to Beta-Lactam Antibiotics. mBio. 2021;12(3):e0053021.

85. Harper L, Balasubramanian D, Ohneck EA, Sause WE, Chapman J, Mejia-Sosa B, et al. Staphylococcus aureus Responds to the Central Metabolite Pyruvate To Regulate Virulence. mBio. 2018;9(1).

86. Tomlinson KL, Riquelme SA. Host-bacteria metabolic crosstalk drives S. aureus biofilm. Microb Cell. 2021;8(5):106–7.

87. Tomlinson KL, Lung TWF, Dach F, Annavajhala MK, Gabryszewski SJ, Groves RA, et al. Staphylococcus aureus induces an itaconate-dominated immunometabolic response that drives biofilm formation. Nat Commun. 2021;12(1):1399.

88. Passalacqua KD, Charbonneau ME, O’Riordan MXD. Bacterial Metabolism Shapes the Host-Pathogen Interface. Microbiol Spectr. 2016;4(3).

89. Zhao R, Xu L, Chen J, Yang Y, Guo X, Dai M, et al. Itaconate induces tolerance of Staphylococcus aureus to aminoglycoside antibiotics. Front Microbiol. 2024;15:1450085.

90. Ramond E, Gesbert G, Rigard M, Dairou J, Dupuis M, Dubail I, et al. Glutamate utilization couples oxidative stress defense and the tricarboxylic acid cycle in Francisella phagosomal escape. PLoS Pathog. 2014;10(1):e1003893.

91. Tavares AF, Teixeira M, Romão CC, Seixas JD, Nobre LS, Saraiva LM. Reactive oxygen species mediate bactericidal killing elicited by carbon monoxide-releasing molecules. J Biol Chem. 2011;286(30):26708–17.

92. Yee R, Cui P, Shi W, Feng J, Zhang Y. Genetic Screen Reveals the Role of Purine Metabolism in Staphylococcus aureus Persistence to Rifampicin. Antibiotics (Basel). 2015;4(4):627–42.

93. Yee R, Feng J, Wang J, Chen J, Zhang Y. Identification of Genes Regulating Cell Death in Staphylococcus aureus. Front Microbiol. 2019;10:2199.

94. Malachowa N, Whitney AR, Kobayashi SD, Sturdevant DE, Kennedy AD, Braughton KR, et al. Global changes in Staphylococcus aureus gene expression in human blood. PLoS One. 2011;6(4):e18617.

95. Haupt K, Reuter M, van den Elsen J, Burman J, Hälbich S, Richter J, et al. The Staphylococcus aureus protein Sbi acts as a complement inhibitor and forms a tripartite complex with host complement Factor H and C3b. PLoS Pathog. 2008;4(12):e1000250.

96. Yan J, Han D, Liu C, Gao Y, Li D, Liu Y, et al. Staphylococcus aureus VraX specifically inhibits the classical pathway of complement by binding to C1q. Mol Immunol. 2017;88:38–44.

97. Spaan AN, Vrieling M, Wallet P, Badiou C, Reyes-Robles T, Ohneck EA, et al. The staphylococcal toxins γ-haemolysin AB and CB differentially target phagocytes by employing specific chemokine receptors. Nat Commun. 2014;5:5438.

98. Zhao Y, van Kessel KPM, de Haas CJC, Rogers MRC, van Strijp JAG, Haas PA. Staphylococcal superantigen-like protein 13 activates neutrophils via formyl peptide receptor 2. Cell Microbiol. 2018;20(11):e12941.

99. Higgins J, Loughman A, van Kessel KP, van Strijp JA, Foster TJ. Clumping factor A of Staphylococcus aureus inhibits phagocytosis by human polymorphonuclear leucocytes. FEMS Microbiol Lett. 2006;258(2):290–6.

100. Smith EJ, Visai L, Kerrigan SW, Speziale P, Foster TJ. The Sbi protein is a multifunctional immune evasion factor of Staphylococcus aureus. Infect Immun. 2011;79(9):3801–9.

101. Chen F, Zhao Q, Yang Z, Chen R, Pan H, Wang Y, et al. Citrate serves as a signal molecule to modulate carbon metabolism and iron homeostasis in Staphylococcus aureus. PLoS Pathog. 2024;20(7):e1012425.

102. Poudel S, Tsunemoto H, Seif Y, Sastry AV, Szubin R, Xu S, et al. Revealing 29 sets of independently modulated genes in Staphylococcus aureus, their regulators, and role in key physiological response. Proc Natl Acad Sci U S A. 2020;117(29):17228–39.

103. Cascales E. Inside the Chamber of Secrets of the Type III Secretion System. Cell. 2017;168(6):949–51.

104. Fey PD, Endres JL, Yajjala VK, Widhelm TJ, Boissy RJ, Bose JL, et al. A genetic resource for rapid and comprehensive phenotype screening of nonessential Staphylococcus aureus genes. mBio. 2013;4(1):e00537–12.

105. Wick RR, Judd LM, Cerdeira LT, Hawkey J, Méric G, Vezina B, et al. Trycycler: consensus long-read assemblies for bacterial genomes. Genome Biol. 2021;22(1):266.

106. Darling AC, Mau B, Blattner FR, Perna NT. Mauve: multiple alignment of conserved genomic sequence with rearrangements. Genome Res. 2004;14(7):1394–403.

