## Supplemental figures for "Integrated multi-omics reveals coordinated *Staphylococcus aureus* metabolic, iron transport, and stress responses to human serum"

### Supplementary Figures

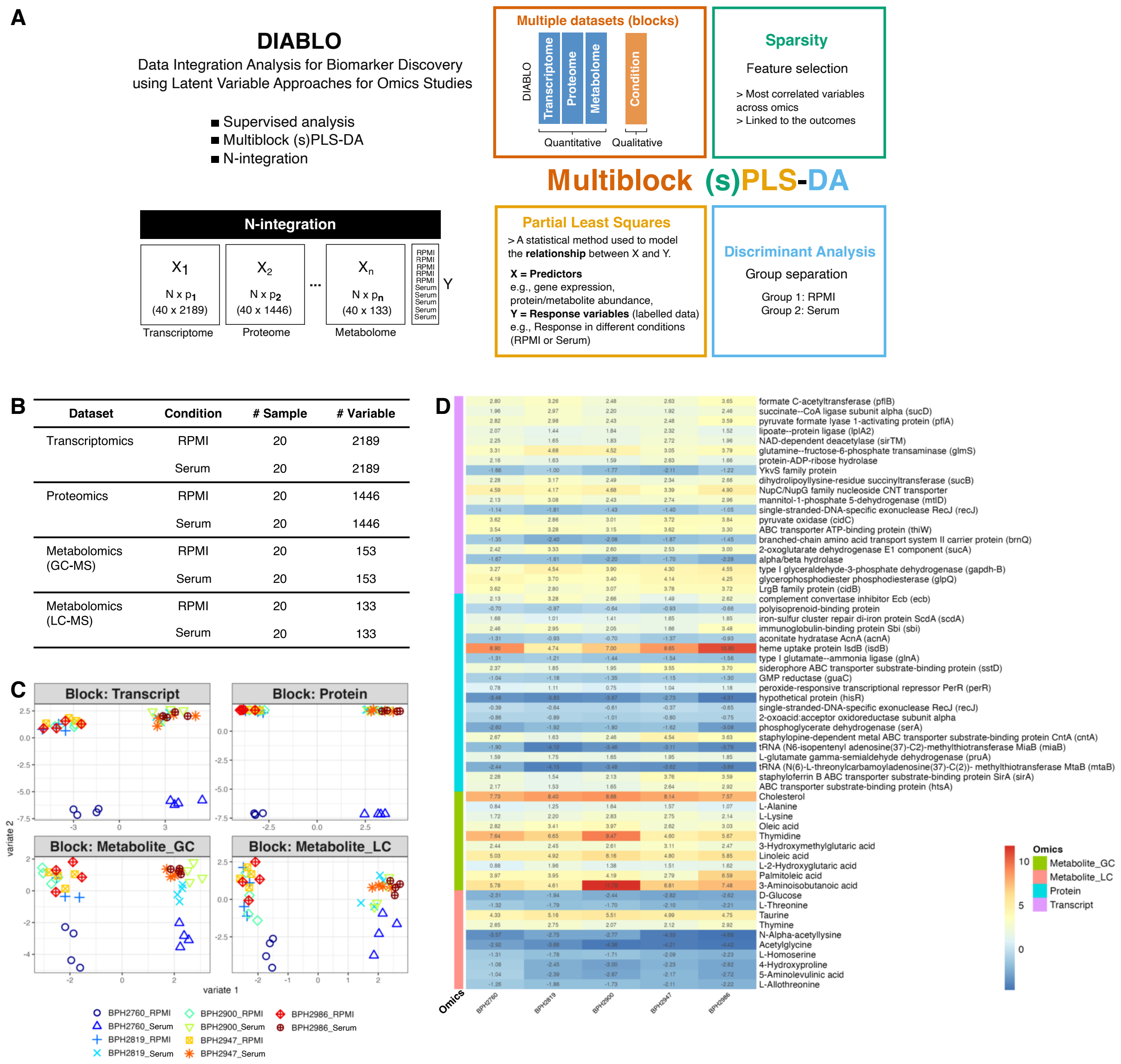


**Figure S1. Multi-Omics variables used for analysis (related to Figure 1)**
(A) Schematic diagram of the DIABLO framework. DIABLO performs a supervised analysis based on a multiblock (s)PLS-DA model (left) and an N-integration strategy across omics datasets (right). (B) Omic datasets used for multi-omics analysis. (C) Sample plot from multiblock sPLS-DA according to their scores on the first 2 components for each dataset. Samples from each dataset (transcriptomics, proteomics, and metabolomics (GC-MS and LC-MS)) are coloured by strains (BPH2760, BPH2819, BPH2900, BPH2947, and BPH2986). (D) Heatmap representing expression or abundance data (log2FC compared to RPMI control) of selected variables.

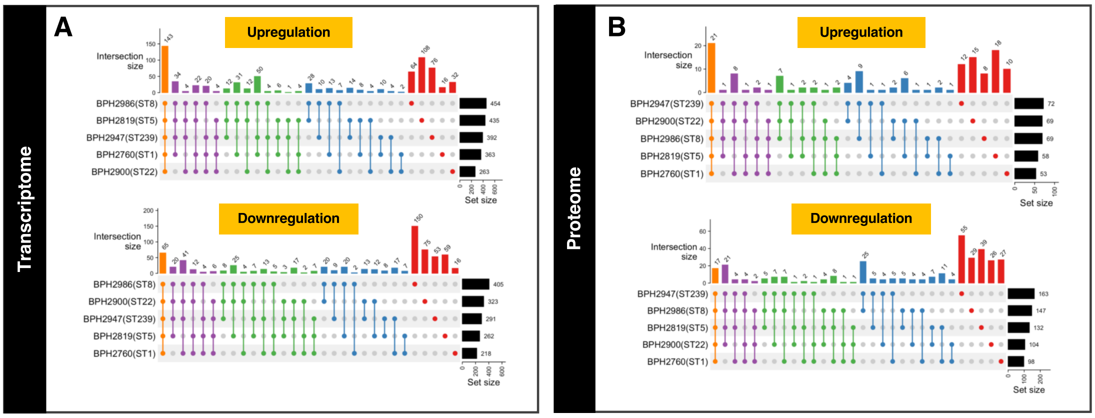


**Figure S2. Differentially Expressed Genes/Proteins of *S. aureus* Exposed to Serum (related to Figure 2)**

Upset plots representing the shared and distinctive differentially expressed (DE) genes/proteins in human serum, across five distinct strains of *S. aureus* (A-B). (A) Up-regulated and down-regulated DE genes. (B) Up-regulated and down-regulated DE proteins. In each panel, the right horizontal bar graph represents the total number of up- or down-regulated DE genes/proteins for individual *S. aureus* strains. The filled circles in each panel's matrix indicate strains that are included in the intersections. Connected circles indicate intersecting DE genes/proteins between strains. The top bar graph in each panel indicates the number of DE genes/proteins for each unique or overlapping combination.

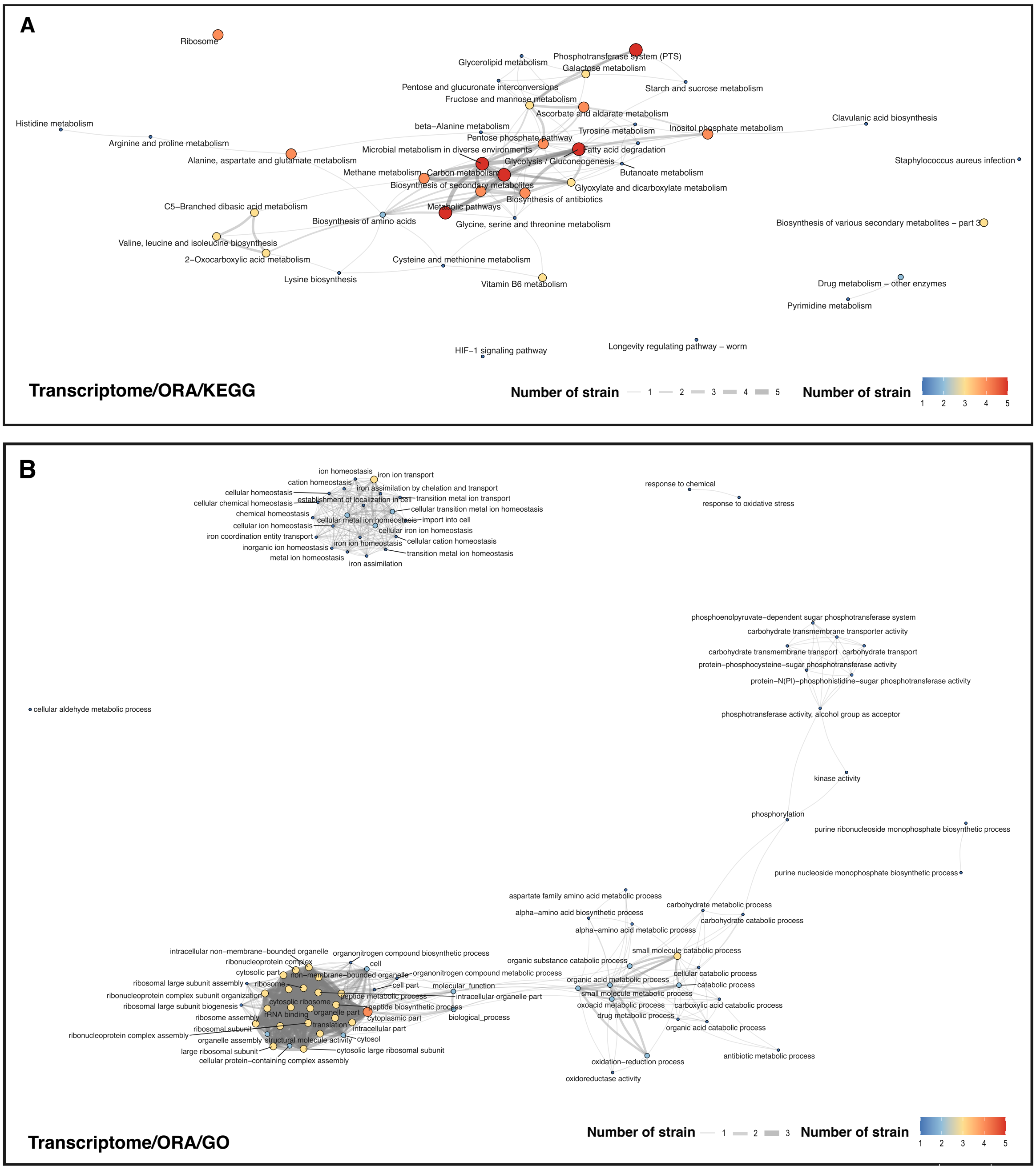
A
Ribosome
Glycerolipid metabolism
Phosphotransferase system (PTS)
Galactose metabolism
Pentose and glucuronate interconversions
Starch and sucrose metabolism
Fructose and mannose metabolism
Ascorbate and aldarate metabolism
Histidine metabolism
beta−Alanine metabolism
Tyrosine metabolism
Inositol phosphate metabolism
Clavulanic acid biosynthesis
Arginine and proline metabolism
Pentose phosphate pathway
Microbial metabolism in diverse environments
Fatty acid degradation
Alanine, aspartate and glutamate metabolism
Glycolysis / Gluconeogenesis
Staphylococcus aureus infection
Methane metabolism
Carbon metabolism
Butanoate metabolism
Biosynthesis of secondary metabolites
Glyoxylate and dicarboxylate metabolism
Biosynthesis of antibiotics
C5−Branched dibasic acid metabolism
Metabolic pathways
Biosynthesis of amino acids
Glycine, serine and threonine metabolism
Biosynthesis of various secondary metabolites – part 3
Valine, leucine and isoleucine biosynthesis
2−Oxocarboxylic acid metabolism
Cysteine and methionine metabolism
Lysine biosynthesis
Vitamin B6 metabolism
Drug metabolism – other enzymes
Pyrimidine metabolism
Longevity regulating pathway – worm
HIF−1 signaling pathway
Transcriptome/ORA/KEGG
Number of strain
1 2 3 4 5
Number of strain
1 2 3 4 5
B
ion homeostasis
iron ion transport
cation homeostasis
iron assimilation by chelation and transport
cellular homeostasis
establishment of localization in cell
transition metal ion transport
cellular chemical homeostasis
cellular transition metal ion homeostasis
chemical homeostasis
cellular metal ion homeostasis
import into cell
cellular ion homeostasis
cellular iron ion homeostasis
iron coordination entity transport
iron ion homeostasis
cellular cation homeostasis
inorganic ion homeostasis
transition metal ion homeostasis
metal ion homeostasis
iron assimilation
response to chemical
response to oxidative stress
phosphoenolpyruvate−dependent sugar phosphotransferase system
carbohydrate transmembrane transporter activity
carbohydrate transmembrane transport
carbohydrate transport
protein−phosphocysteine−sugar phosphotransferase activity
protein−N(PI)−phosphohistidine−sugar phosphotransferase activity
cellular aldehyde metabolic process
phosphotransferase activity, alcohol group as acceptor
kinase activity
phosphorylation
purine ribonucleoside monophosphate biosynthetic process
purine nucleoside monophosphate biosynthetic process
aspartate family amino acid metabolic process
carbohydrate metabolic process
alpha−amino acid biosynthetic process
carbohydrate catabolic process
alpha−amino acid metabolic process
intracellular non−membrane−bounded organelle
organonitrogen compound biosynthetic process
small molecule catabolic process
ribonucleoprotein complex
cell
organic substance catabolic process
cytosolic part
non−membrane−bounded organelle
organonitrogen compound metabolic process
cellular catabolic process
ribosomal large subunit assembly
ribosome
cell part
molecular_function
organic acid metabolic process
catabolic process
ribonucleoprotein complex subunit organization
peptide metabolic process
intracellular organelle part
small molecule metabolic process
ribosomal large subunit biogenesis
cytosolic ribosome
peptide biosynthetic process
oxoacid metabolic process
carboxylic acid catabolic process
ribosome assembly
rRNA binding
organelle part
cytoplasmic part
biological_process
drug metabolic process
ribosomal subunit
translation
intracellular part
organic acid catabolic process
ribonucleoprotein complex assembly
organelle assembly
structural molecule activity
cytosol
large ribosomal subunit
cytosolic large ribosomal subunit
antibiotic metabolic process
cellular protein−containing complex assembly
oxidation−reduction process
oxidoreductase activity
Transcriptome/ORA/GO
Number of strain
1 2 3
Number of strain
1 2 3 4 5

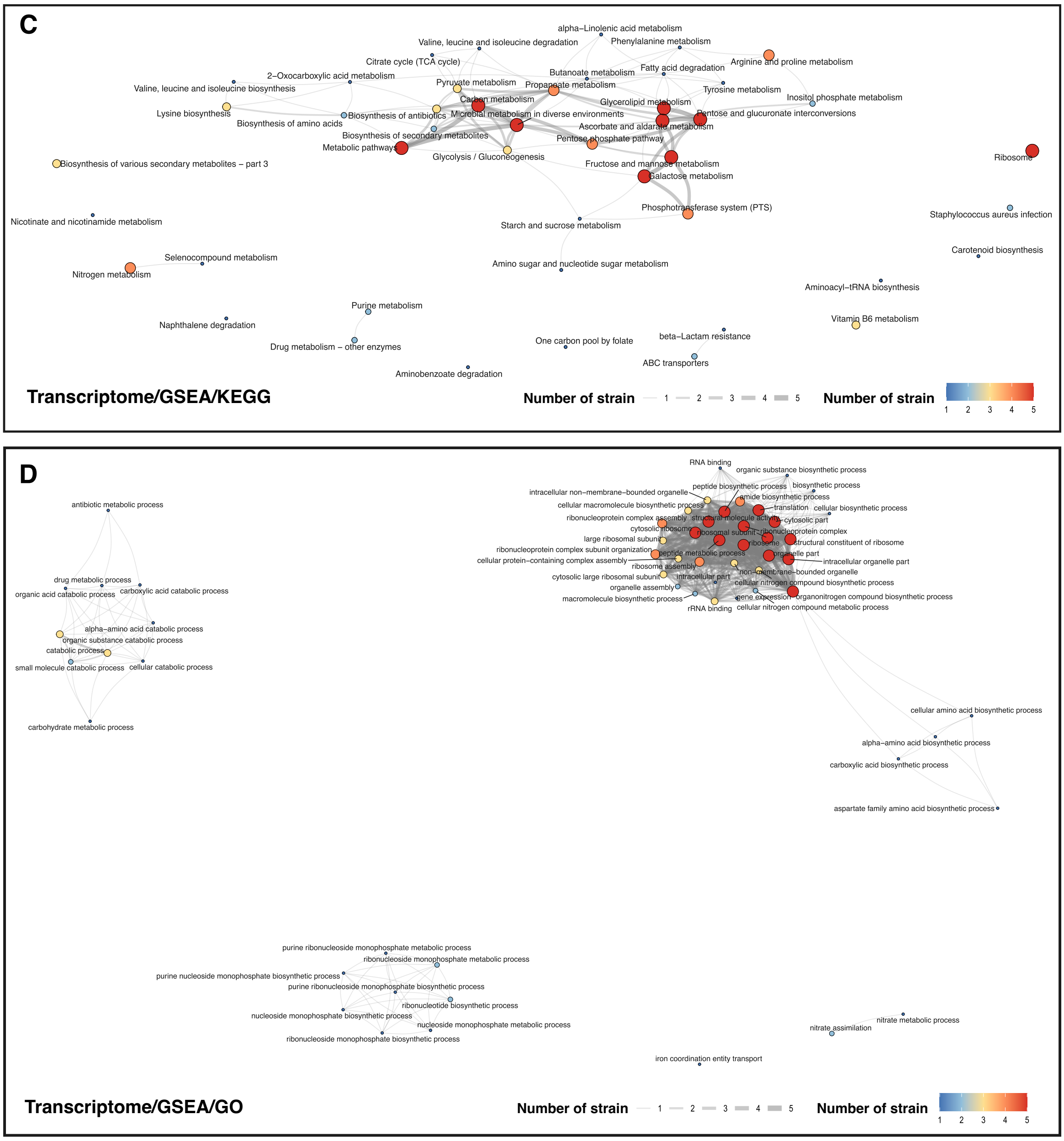
C
alpha–Linolenic acid metabolism
Valine, leucine and isoleucine degradation
Phenylalanine metabolism
Citrate cycle (TCA cycle)
Arginine and proline metabolism
Fatty acid degradation
2–Oxocarboxylic acid metabolism
Butanoate metabolism
Valine, leucine and isoleucine biosynthesis
Pyruvate metabolism
Propanoate metabolism
Tyrosine metabolism
Inositol phosphate metabolism
Carbon metabolism
Glycerolipid metabolism
Lysine biosynthesis
Biosynthesis of antibiotics
Microbial metabolism in diverse environments
Pentose and glucuronate interconversions
Biosynthesis of amino acids
Ascorbate and aldarate metabolism
Biosynthesis of secondary metabolites
Pentose phosphate pathway
Metabolic pathways
Glycolysis / Gluconeogenesis
Ribosome
Biosynthesis of various secondary metabolites – part 3
Fructose and mannose metabolism
Galactose metabolism
Phosphotransferase system (PTS)
Staphylococcus aureus infection
Nicotinate and nicotinamide metabolism
Starch and sucrose metabolism
Carotenoid biosynthesis
Selenocompound metabolism
Amino sugar and nucleotide sugar metabolism
Nitrogen metabolism
Aminoacyl–tRNA biosynthesis
Purine metabolism
Vitamin B6 metabolism
Naphthalene degradation
beta–Lactam resistance
One carbon pool by folate
Drug metabolism – other enzymes
ABC transporters
Aminobenzoate degradation
Transcriptome/GSEA/KEGG
Number of strain
1 2 3 4 5
Number of strain
1 2 3 4 5
D
antibiotic metabolic process
drug metabolic process
organic acid catabolic process
carboxylic acid catabolic process
alpha–amino acid catabolic process
organic substance catabolic process
catabolic process
small molecule catabolic process
cellular catabolic process
carbohydrate metabolic process
RNA binding
organic substance biosynthetic process
peptide biosynthetic process
biosynthetic process
intracellular non–membrane–bounded organelle
amide biosynthetic process
cellular macromolecule biosynthetic process
translation
cellular biosynthetic process
ribonucleoprotein complex assembly
structural molecule activity
cytosolic part
cytosolic ribosome
ribosomal subunit
ribonucleoprotein complex
large ribosomal subunit
ribosome
structural constituent of ribosome
ribonucleoprotein complex subunit organization
peptide metabolic process
organelle part
cellular protein–containing complex assembly
intracellular organelle part
ribosome assembly
non–membrane–bounded organelle
cytosolic large ribosomal subunit
intracellular part
cellular nitrogen compound biosynthetic process
organelle assembly
gene expression
organonitrogen compound biosynthetic process
macromolecule biosynthetic process
rRNA binding
cellular nitrogen compound metabolic process
cellular amino acid biosynthetic process
alpha–amino acid biosynthetic process
carboxylic acid biosynthetic process
aspartate family amino acid biosynthetic process
purine ribonucleoside monophosphate metabolic process
ribonucleoside monophosphate metabolic process
purine nucleoside monophosphate biosynthetic process
purine ribonucleoside monophosphate biosynthetic process
ribonucleotide biosynthetic process
nucleoside monophosphate biosynthetic process
nucleoside monophosphate metabolic process
ribonucleoside monophosphate biosynthetic process
nitrate metabolic process
nitrate assimilation
iron coordination entity transport
Transcriptome/GSEA/GO
Number of strain
1 2 3 4 5
Number of strain
1 2 3 4 5

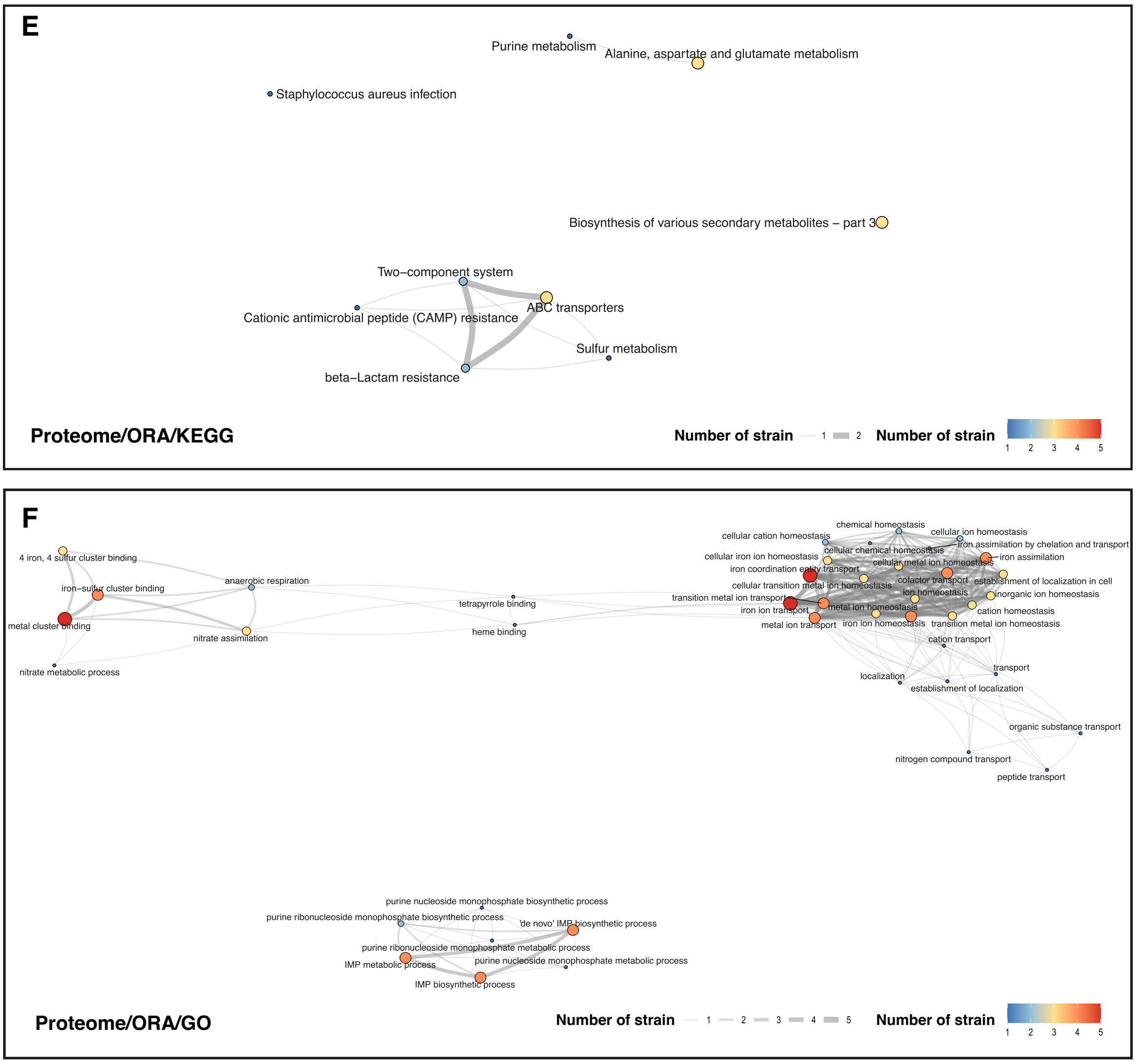
E
Purine metabolism
Alanine, aspartate and glutamate metabolism
Staphylococcus aureus infection
Biosynthesis of various secondary metabolites – part 3
Two–component system
ABC transporters
Cationic antimicrobial peptide (CAMP) resistance
Sulfur metabolism
beta–Lactam resistance
Proteome/ORA/KEGG
Number of strain
1
2
Number of strain
1 2 3 4 5
F
4 iron, 4 sulfur cluster binding
iron–sulfur cluster binding
anaerobic respiration
metal cluster binding
nitrate assimilation
nitrate metabolic process
tetrapyrrole binding
heme binding
chemical homeostasis
cellular cation homeostasis
cellular ion homeostasis
iron assimilation by chelation and transport
cellular iron ion homeostasis
cellular chemical homeostasis
iron assimilation
iron coordination entity transport
cellular metal ion homeostasis
cellular transition metal ion homeostasis
cofactor transport
establishment of localization in cell
transition metal ion transport
ion homeostasis
inorganic ion homeostasis
iron ion transport
metal ion homeostasis
cation homeostasis
metal ion transport
iron ion homeostasis
transition metal ion homeostasis
cation transport
transport
localization
establishment of localization
organic substance transport
nitrogen compound transport
peptide transport
purine nucleoside monophosphate biosynthetic process
purine ribonucleoside monophosphate biosynthetic process
'de novo' IMP biosynthetic process
purine ribonucleoside monophosphate metabolic process
IMP metabolic process
purine nucleoside monophosphate metabolic process
IMP biosynthetic process
Proteome/ORA/GO
Number of strain
1 2 3 4 5
Number of strain
1 2 3 4 5

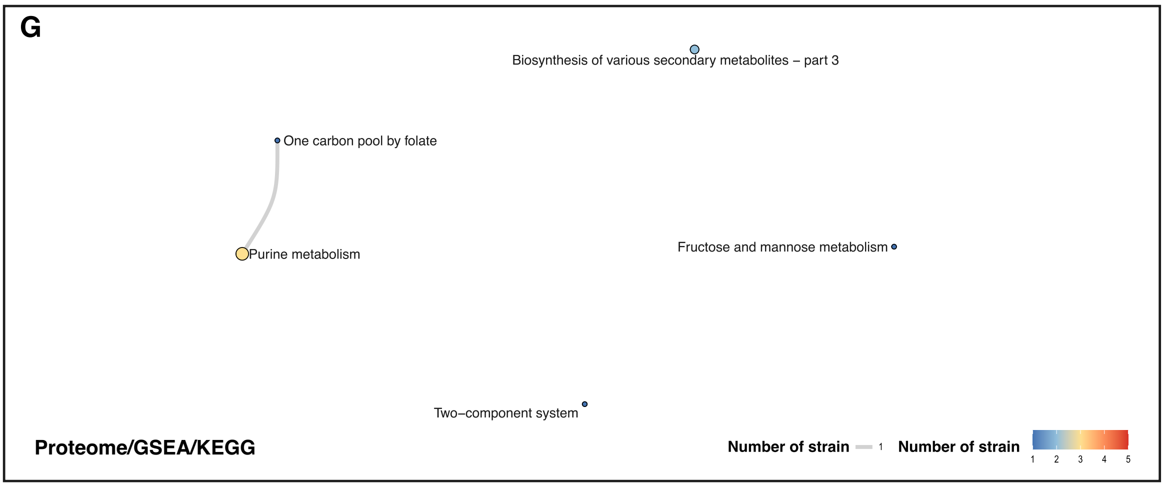

G
Biosynthesis of various secondary metabolites – part 3
One carbon pool by folate
Purine metabolism
Fructose and mannose metabolism
Two–component system
Proteome/GSEA/KEGG
Number of strain
1
Number of strain
1
2
3
4
5


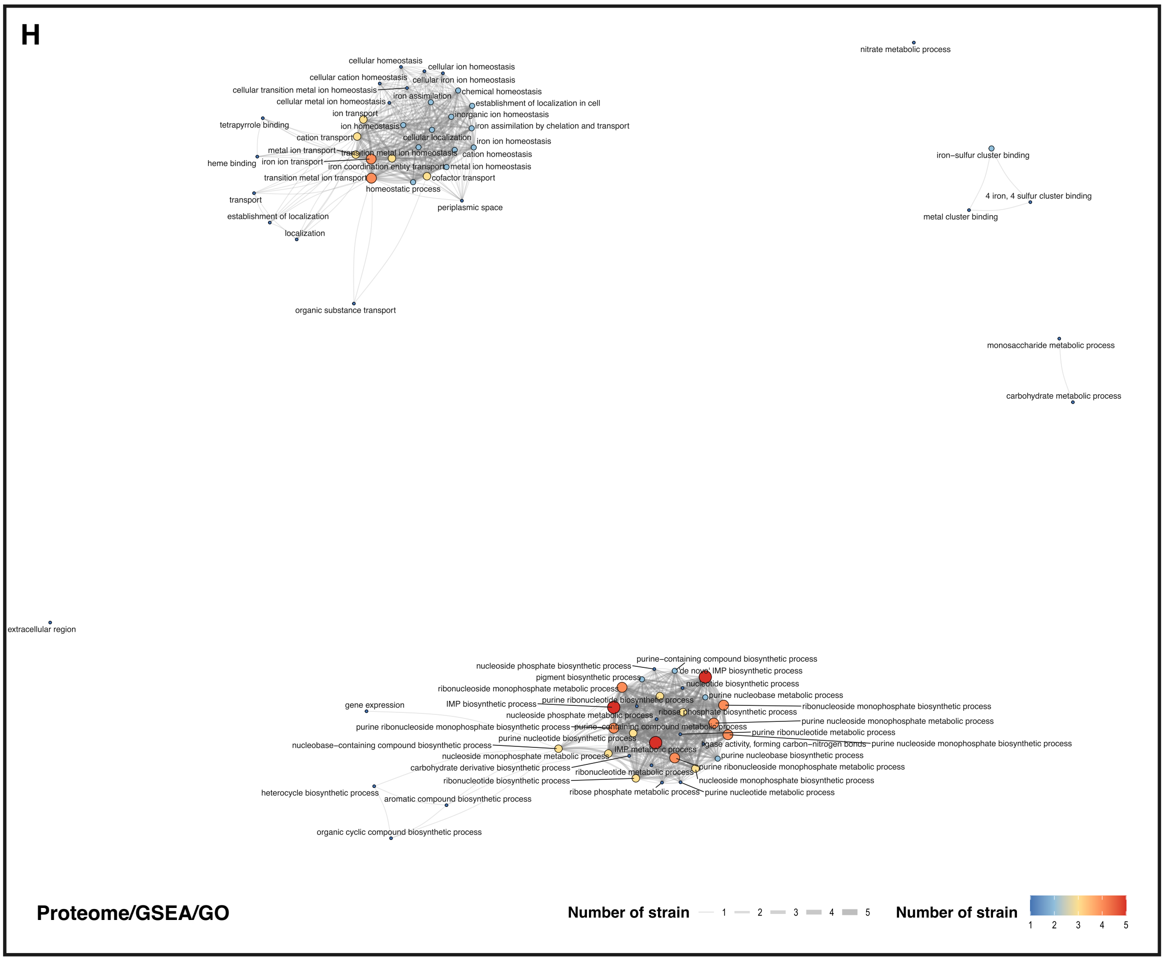

H
cellular homeostasis
cellular ion homeostasis
cellular cation homeostasis
cellular iron ion homeostasis
cellular transition metal ion homeostasis
iron assimilation
chemical homeostasis
cellular metal ion homeostasis
establishment of localization in cell
ion transport
inorganic ion homeostasis
tetrapyrrole binding
ion homeostasis
iron assimilation by chelation and transport
cation transport
cellular localization
iron ion homeostasis
metal ion transport
transition metal ion homeostasis
cation homeostasis
heme binding
iron ion transport
iron coordination entity transport
metal ion homeostasis
transition metal ion transport
cofactor transport
homeostatic process
transport
periplasmic space
establishment of localization
localization
organic substance transport
nitrate metabolic process
iron–sulfur cluster binding
4 iron, 4 sulfur cluster binding
metal cluster binding
monosaccharide metabolic process
carbohydrate metabolic process
extracellular region
purine–containing compound biosynthetic process
nucleoside phosphate biosynthetic process
de novo IMP biosynthetic process
pigment biosynthetic process
nucleotide biosynthetic process
ribonucleoside monophosphate metabolic process
purine nucleobase metabolic process
gene expression
IMP biosynthetic process
purine ribonucleotide biosynthetic process
ribonucleoside monophosphate biosynthetic process
nucleoside phosphate metabolic process
ribose phosphate biosynthetic process
purine ribonucleoside monophosphate biosynthetic process
purine–containing compound metabolic process
purine nucleoside monophosphate metabolic process
purine nucleotide biosynthetic process
purine ribonucleotide metabolic process
nucleobase–containing compound biosynthetic process
IMP metabolic process
ligase activity, forming carbon–nitrogen bonds
purine nucleoside monophosphate biosynthetic process
nucleoside monophosphate metabolic process
purine nucleobase biosynthetic process
carbohydrate derivative biosynthetic process
ribonucleotide metabolic process
purine ribonucleoside monophosphate metabolic process
ribonucleotide biosynthetic process
nucleoside monophosphate biosynthetic process
heterocycle biosynthetic process
ribose phosphate metabolic process
purine nucleotide metabolic process
aromatic compound biosynthetic process
organic cyclic compound biosynthetic process
Proteome/GSEA/GO
Number of strain
1
2
3
4
5
Number of strain
1
2
3
4
5

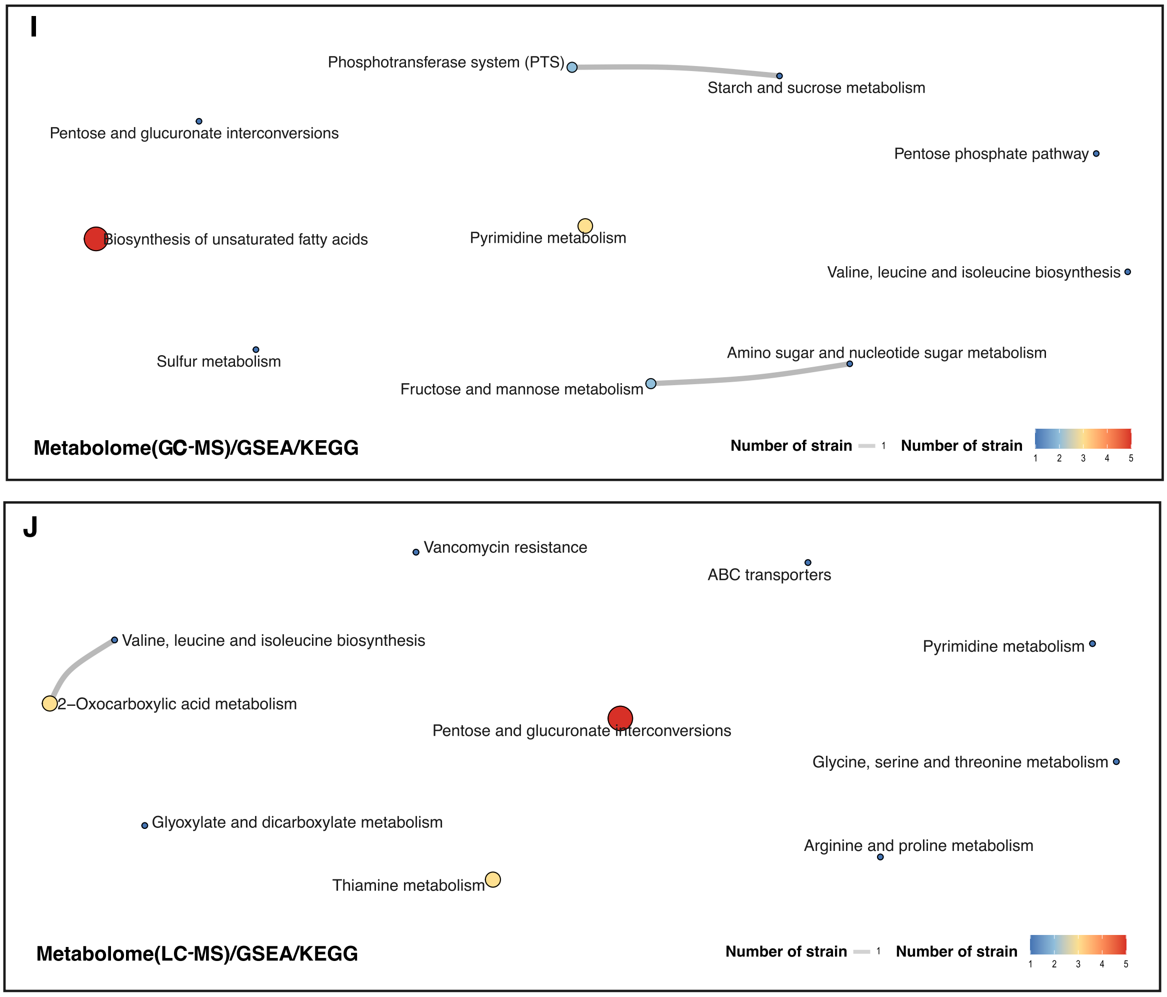


**Figure S3. Extended Network Analyses (related to Figure 2)**
These network analyses use the similarity of gene subsets shared between enriched pathways, obtained from ORA or GSEA, to illustrate their interactions and to identify conserved or strain-specific responses to human serum.
(A-B) ORA and (C-D) GSEA performed on transcriptomic datasets.
(E-F) ORA and (G-H) GSEA performed on proteomic datasets.
(I) GSEA performed on metabolomic (GC-MS) datasets.
(J) GSEA performed on metabolomic (LC-MS) datasets.
The dots' size and colour represent the number of strains sharing the same enriched pathways.
The edges' width indicates the number of strains exhibiting interactions between pathways.

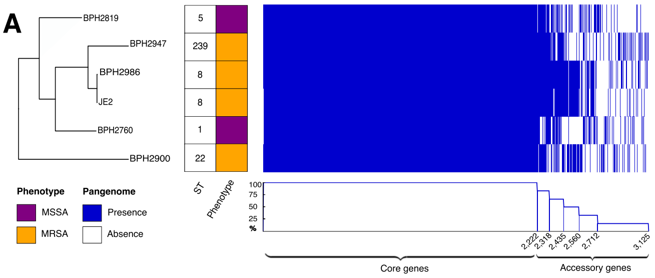


**B**

| Isolate | ST | Allele 1 | Allele 2 | Allele 3 | Allele 4 | Allele 5 | Allele 6 | Allele 7 |
|---|---|---|---|---|---|---|---|---|
| BPH2760 | 1 | arcC(1) | aroE(1) | glpF(1) | gmk(1) | pta(1) | tpi(1) | yqiL(1) |
| BPH2819 | 5 | arcC(1) | aroE(4) | glpF(1) | gmk(4) | pta(12) | tpi(1) | yqiL(10) |
| BPH2900 | 22 | arcC(7) | aroE(6) | glpF(1) | gmk(5) | pta(8) | tpi(8) | yqiL(6) |
| BPH2947 | 239 | arcC(2) | aroE(3) | glpF(1) | gmk(1) | pta(4) | tpi(4) | yqiL(3) |
| BPH2986 | 8 | arcC(3) | aroE(3) | glpF(1) | gmk(1) | pta(4) | tpi(4) | yqiL(3) |
| JE2 | 8 | arcC(3) | aroE(3) | glpF(1) | gmk(1) | pta(4) | tpi(4) | yqiL(3) |

**C**

| Isolate | BPH2760 | BPH2819 | BPH2900 | BPH2947 | BPH2986 | JE2 |
|---|---|---|---|---|---|---|
| BPH2760 | 0 | 17539 | 31553 | 20999 | 15398 | 15355 |
| BPH2819 | 17539 | 0 | 32163 | 22677 | 17382 | 17341 |
| BPH2900 | 31553 | 32163 | 0 | 34352 | 31989 | 31950 |
| BPH2947 | 20999 | 22677 | 34352 | 0 | 9368 | 9325 |
| BPH2986 | 15398 | 17382 | 31989 | 9368 | 0 | 73 |
| JE2 | 15355 | 17341 | 31950 | 9325 | 73 | 0 |

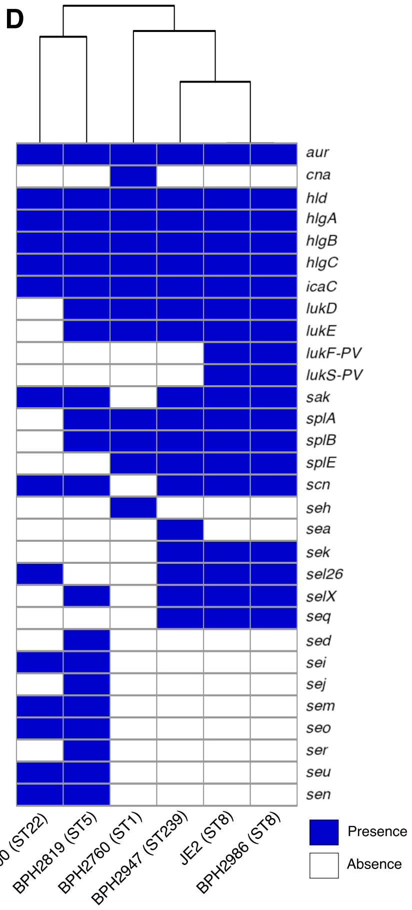


**Figure S4. Genome and Pangenome Analyses of Studied *S. aureus* Strains**
(A) Phylogenetic tree constructed from core genome SNP profiling, accompanied by ST, methicillin resistance phenotype, and pangenome content. (B) *In silico* MLST of studied *S. aureus* strains performed using the comparison of seven housekeeping genes. (C) Core genome SNP profiles for generation of the phylogenetic tree in Figure S4A. (D) Heatmap representing the presence of virulence genes.

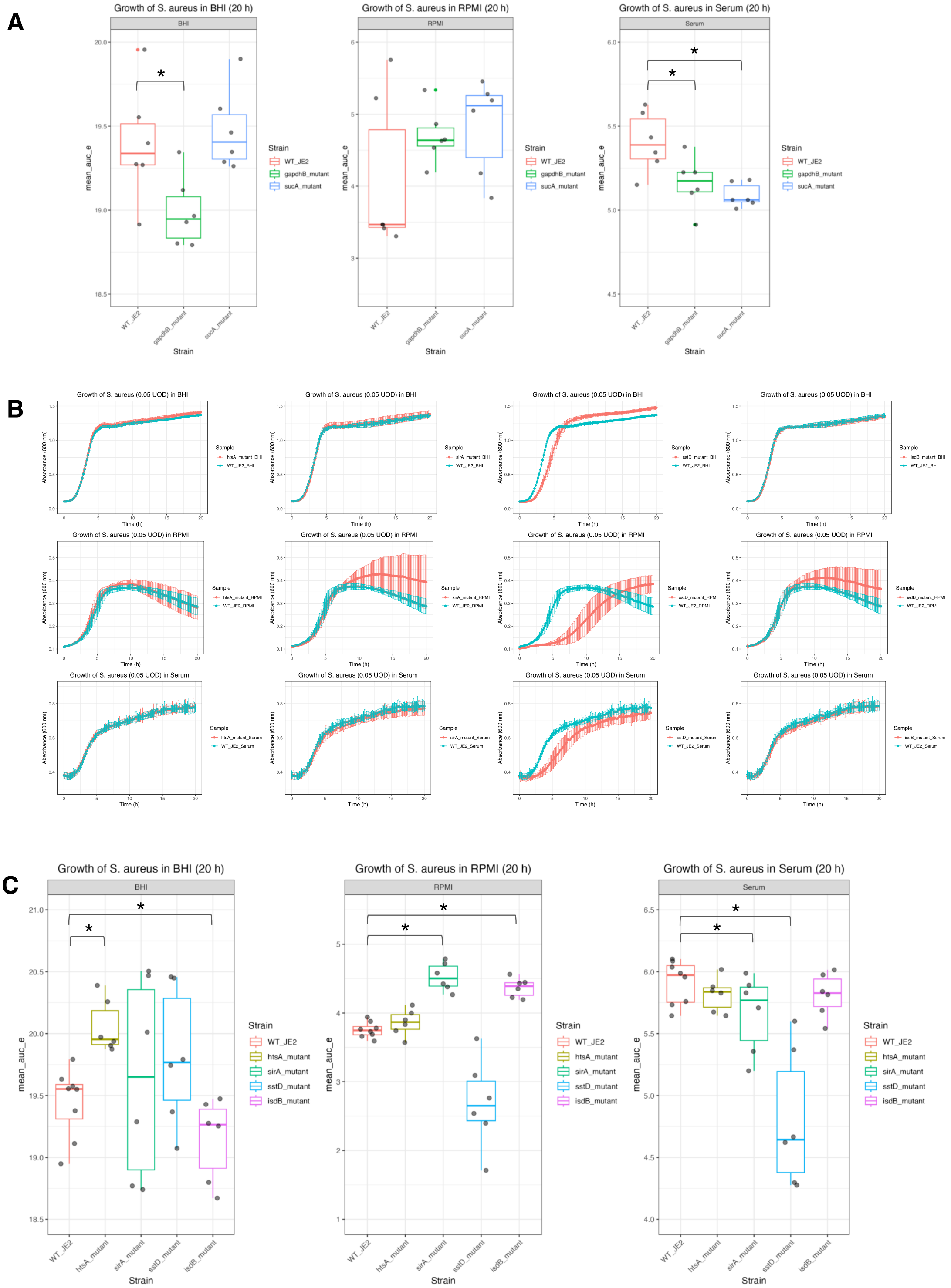
A
Growth of S. aureus in BHI (20 h)
BHI
Growth of S. aureus in RPMI (20 h)
RPMI
Growth of S. aureus in Serum (20 h)
Serum
mean_auc_e
Strain
WT_JE2
gapdhB_mutant
sucA_mutant
*
B
Growth of S. aureus (0.05 UOD) in BHI
Growth of S. aureus (0.05 UOD) in RPMI
Growth of S. aureus (0.05 UOD) in Serum
Absorbance (600 nm)
Time (h)
Sample
C
Growth of S. aureus in BHI (20 h)
BHI
Growth of S. aureus in RPMI (20 h)
RPMI
Growth of S. aureus in Serum (20 h)
Serum
mean_auc_e
Strain
WT_JE2
htsA_mutant
sirA_mutant
sstD_mutant
isdB_mutant
*

**D**

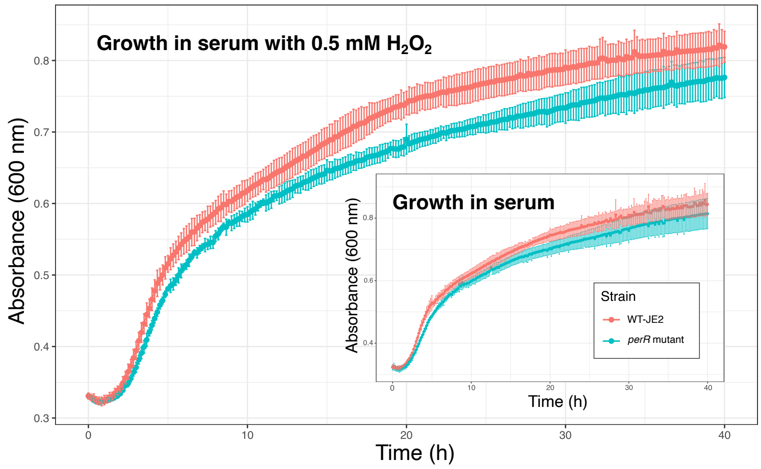


**E**

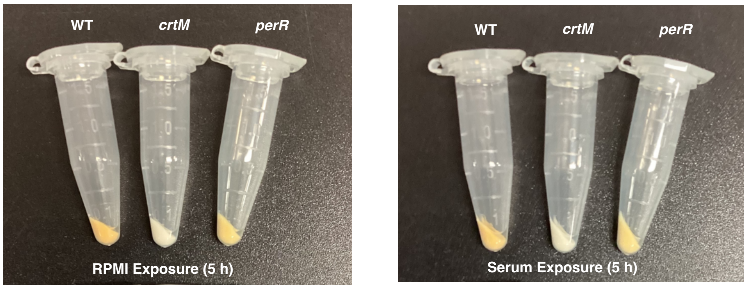


**Figure S5. *S. aureus* Fitness in Human Serum and Staphyloxanthin Production (related to Figure 3)**

(A) Box plots comparing Area-under-the-curve (AUC) values of *gapdhB* and *sucA* mutants to wild-type *S. aureus* (WT-JE2) cultured in BHI, RPMI and serum for 20 hours. (B) Representative growth curves for the *S. aureus* WT- JE2 and iron transport-deficient mutants (*htsA*, *sirA*, *sstD*, *isdB*) cultured in BHI, RPMI and serum for 20 hours. (C) Box plots comparing AUC values of iron transport-deficient mutants (*htsA*, *sirA*, *sstD*, *isdB*) with WT-JE2 cultured in serum for 20 hours. (D) Representative growth curves for the *S. aureus* WT-JE2 and *perR* mutant grown in serum alone (inset plot) and serum containing $H_2O_2$ (large plot) over 40 hours. (E) Pigmentation phenotype of WT-JE2, *crtM* mutant and *perR* mutant after 5 h incubation in RPMI (left) or serum (right) .

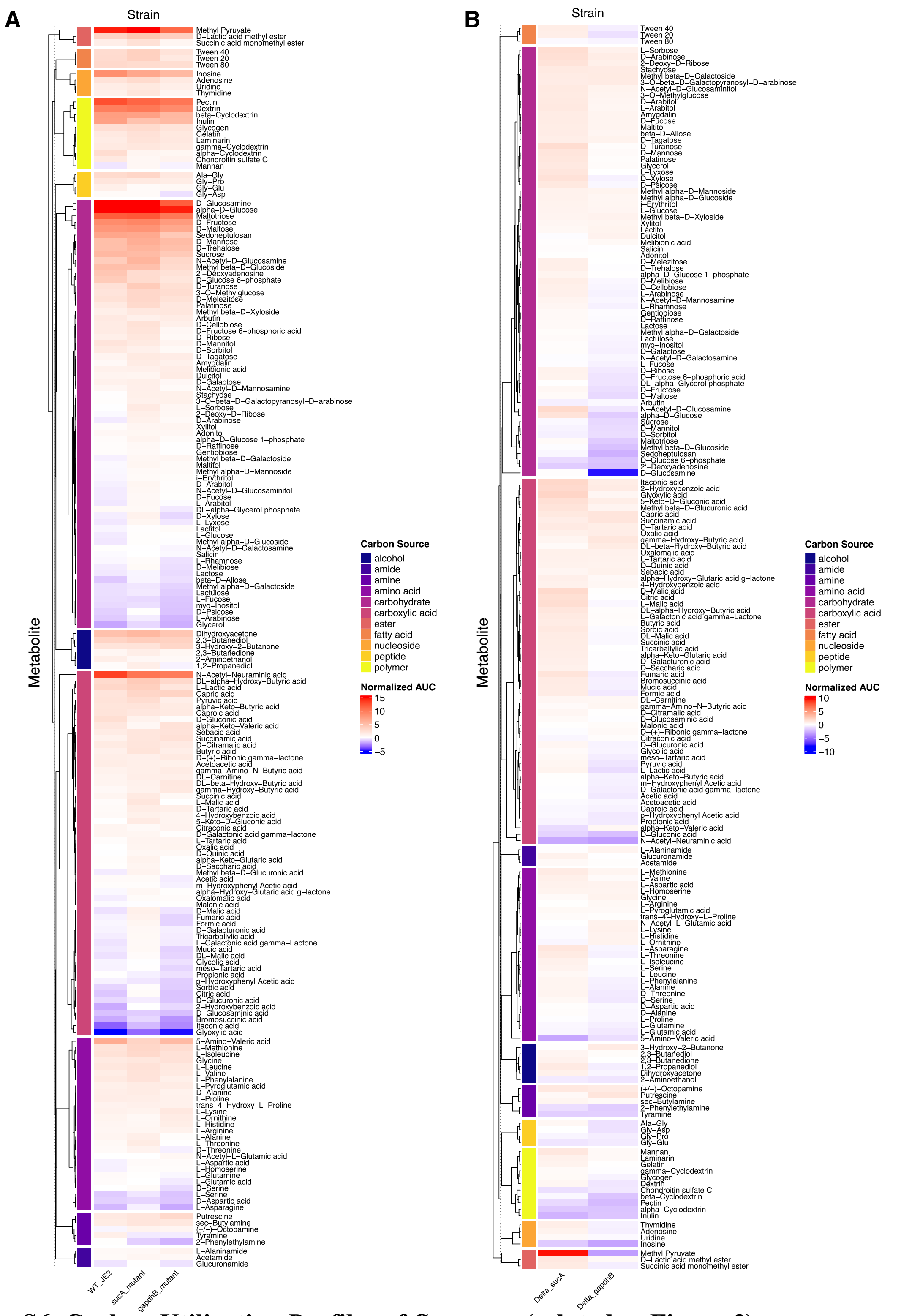


**Figure S6. Carbon Utilization Profiles of *S. aureus* (related to Figure 3)**

(A) Heatmap representing carbon utilization profiles of WT-JE2, *gapdhB* mutant, and *sucA* mutant. Metabolites are clustered based on carbon source types, following substrate classification (Biolog). The normalized area under the curve (AUC) for each metabolite was calculated and subtracted from the AUC of the negative control to represent utilization levels.

(B) Heatmap showing the differences in metabolic activity between the mutants (*gapdhB* and *sucA*) and WT-JE2. The normalized AUC values represent the delta between *sucA* *gapdhB* mutant and WT-JE2, highlighting relative changes in metabolic utilization.

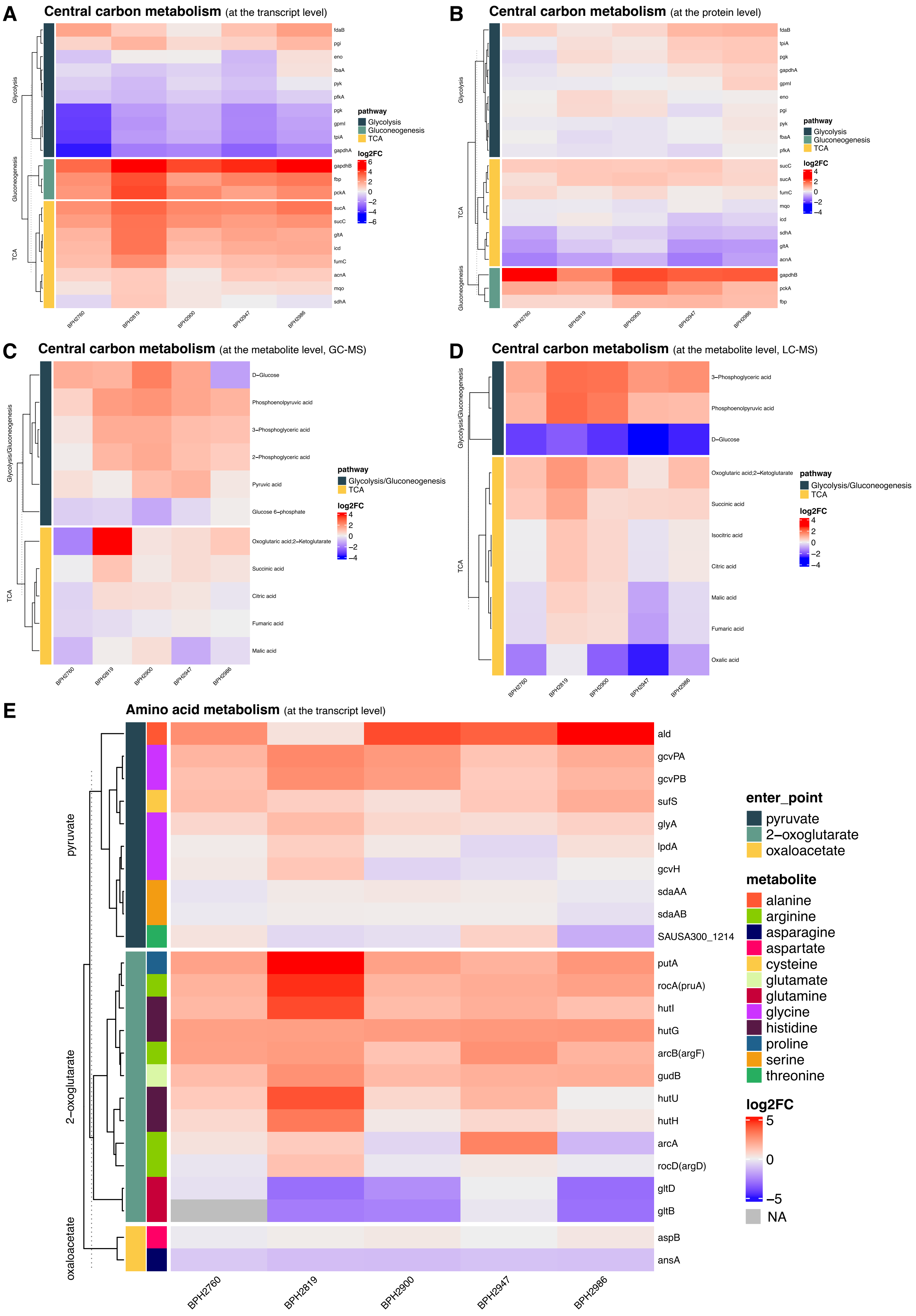
A
Central carbon metabolism (at the transcript level)
Glycolysis
Gluconeogenesis
TCA
fdaB
pgi
eno
fbaA
pyk
pfkA
pgk
gpmI
tpiA
gapdhA
gapdhB
fbp
pckA
sucA
sucC
gltA
icd
fumC
acnA
mqo
sdhA
BPH2760
BPH2819
BPH2900
BPH2947
BPH2986
pathway
Glycolysis
Gluconeogenesis
TCA
log2FC
6
4
2
0
−2
−4
−6
B
Central carbon metabolism (at the protein level)
Glycolysis
TCA
Gluconeogenesis
fdaB
tpiA
pgk
gapdhA
gpmI
eno
pgi
pyk
fbaA
pfkA
sucC
sucA
fumC
mqo
icd
sdhA
gltA
acnA
gapdhB
pckA
fbp
BPH2760
BPH2819
BPH2900
BPH2947
BPH2986
pathway
Glycolysis
Gluconeogenesis
TCA
log2FC
4
2
0
−2
−4
C
Central carbon metabolism (at the metabolite level, GC-MS)
Glycolysis/Gluconeogenesis
TCA
D−Glucose
Phosphoenolpyruvic acid
3−Phosphoglyceric acid
2−Phosphoglyceric acid
Pyruvic acid
Glucose 6−phosphate
Oxoglutaric acid;2−Ketoglutarate
Succinic acid
Citric acid
Fumaric acid
Malic acid
BPH2760
BPH2819
BPH2900
BPH2947
BPH2986
pathway
Glycolysis/Gluconeogenesis
TCA
log2FC
4
2
0
−2
−4
D
Central carbon metabolism (at the metabolite level, LC-MS)
Glycolysis/Gluconeogenesis
TCA
3−Phosphoglyceric acid
Phosphoenolpyruvic acid
D−Glucose
Oxoglutaric acid;2−Ketoglutarate
Succinic acid
Isocitric acid
Citric acid
Malic acid
Fumaric acid
Oxalic acid
BPH2760
BPH2819
BPH2900
BPH2947
BPH2986
pathway
Glycolysis/Gluconeogenesis
TCA
log2FC
4
2
0
−2
−4
E
Amino acid metabolism (at the transcript level)
pyruvate
2−oxoglutarate
oxaloacetate
ald
gcvPA
gcvPB
sufS
glyA
lpdA
gcvH
sdaAA
sdaAB
SAUSA300_1214
putA
rocA(pruA)
hutI
hutG
arcB(argF)
gudB
hutU
hutH
arcA
rocD(argD)
gltD
gltB
aspB
ansA
BPH2760
BPH2819
BPH2900
BPH2947
BPH2986
enter_point
pyruvate
2−oxoglutarate
oxaloacetate
metabolite
alanine
arginine
asparagine
aspartate
cysteine
glutamate
glutamine
glycine
histidine
proline
serine
threonine
log2FC
5
0
−5
NA

**Figure S7. Alterations in Central Carbon Metabolism of *S. aureus* Exposed to Human Serum (related to Figure 4)**

(A-B) Gene expression profiles of *S. aureus* genes involved in glycolysis, gluconeogenesis, and the TCA cycle following human serum exposure, presented at the transcript level (A) and protein level (B). (C-D) Abundance of metabolites detected from GC/MS (C) and LC/MS (D) methods associated with central carbon metabolism following serum exposure. (E) Gene expression profiles of *S. aureus* enzymes responsible for amino acid utilization in the serum environment.

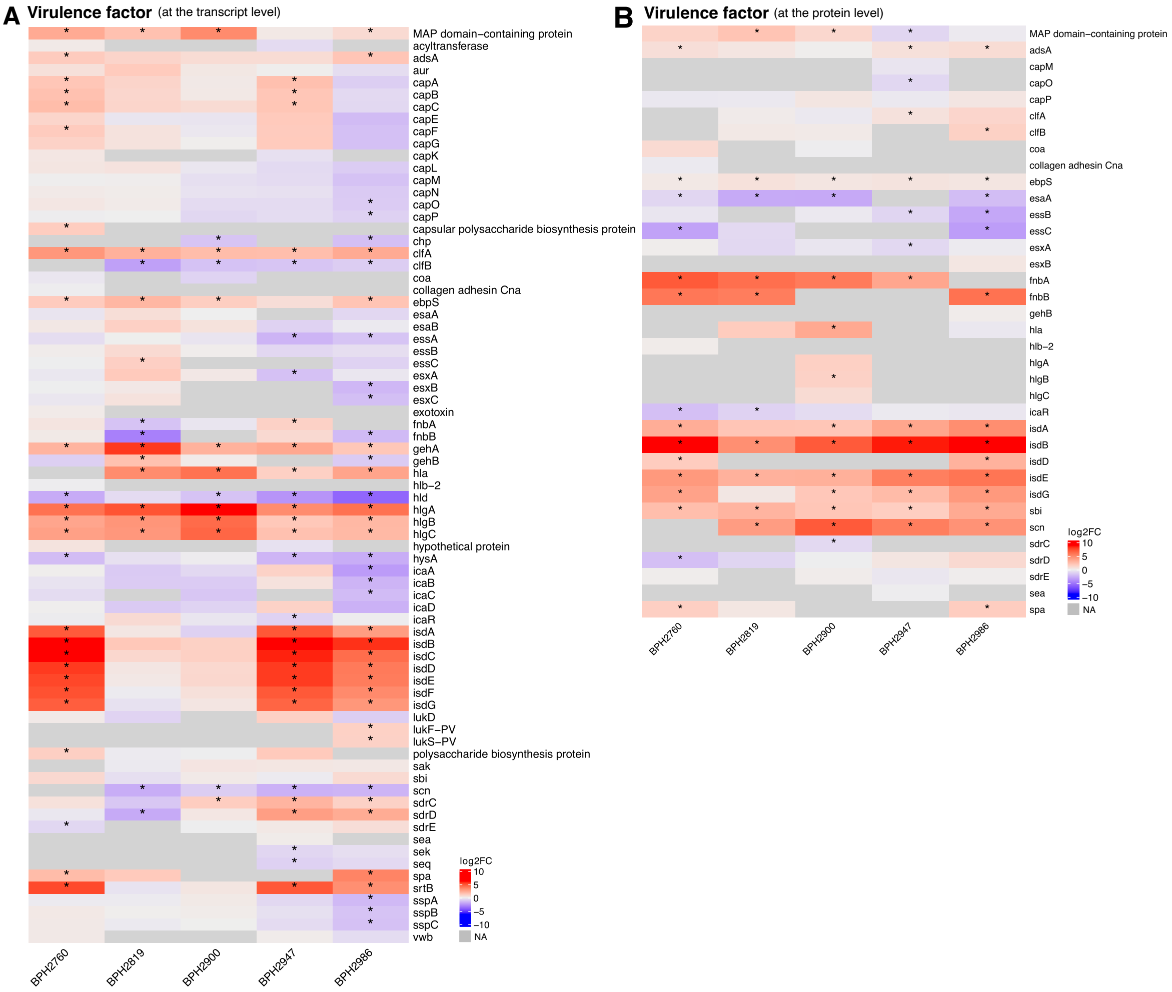


**Figure S8. Alterations in Virulence Profiles of *S. aureus* Exposed to Human Serum**
Gene expression profiles of *S. aureus* virulence factors under human serum exposure, presented at the transcript level (A) and protein level (B).

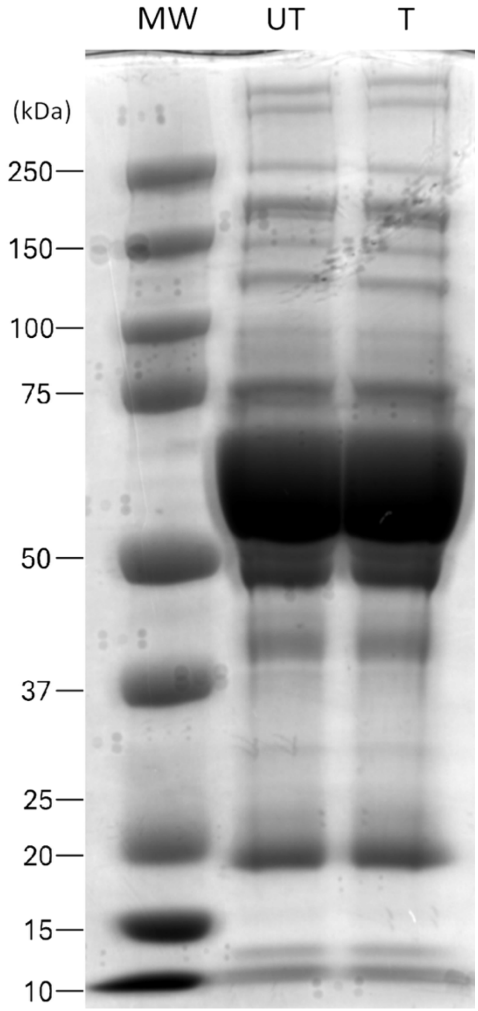


**Figure S9.** Proteins of human serum, prior (UT) and after (T) centrifugation, were analyzed by SDS-PAGE separation and stained with Coomassie blue. MW= Molecular weights.
